# A Novel Biofilm Amyloid Ligand for Caspase-11 Drives Hyperactivation in Dendritic Cells

**DOI:** 10.64898/2026.09.29.755159

**Authors:** Shrutika Mintri, Guangnan Hu, Anukriti Mathur, Leah M. Whiteman, Alexandre Dinis, Brian Le, Kaitlyn Grando, Patience Shumba, Vijay A. Rathinam, Egil Lien, Çağla Tükel, Katherine A. Fitzgerald, Stefania Gallucci

**Affiliations:** Laboratory of Dendritic Cell Biology, Division of Innate Immunity, Department of Medicine, University of Massachusetts Chan Medical School, Worcester, MA, 01545, USA; Division of Innate Immunity, Department of Medicine, University of Massachusetts Chan Medical School, Worcester, MA, 01545, USA; Division of Infectious Diseases and Immunology, Department of Medicine, University of Massachusetts Chan Medical School, Worcester, MA, 01545, USA; Center for Microbiology and Immunology, Lewis Katz School of Medicine, Temple University, Philadelphia, PA, 19140, USA; Department of Immunology, University of Connecticut Health School of Medicine, Farmington, CT, 06030, USA; Center for Molecular inflammation Research, Norwegian University of Science and Technology (NTNU), 7491 Trondheim, Norway

## Abstract

Bacterial biofilms drive persistent infection and chronic inflammation, yet host sensing of biofilm-derived ligands remains poorly understood. Here we identify curli amyloid, an integral component of *Enterobacteriaceae* biofilms, as a novel activator of the non-canonical inflammasome in human and mouse dendritic cells (DCs). Curli both primes and activates the inflammasome, directly binding caspase-11 to engage NLRP3 and drive robust IL-1β maturation without pyroptosis, establishing curli as a previously unrecognized trigger of DC hyperactivation. Mechanistically, this non-lytic signaling is marked by limited gasdermin D oligomerization without mitochondrial localization and NINJ1-dependent membrane rupture. Functionally, curli-hyperactivated DCs show enhanced migration to the draining lymph nodes *in vivo*, suggesting consequent augmented adaptive immunity. Finally, curli-containing biofilms also elicit IL-1β secretion without cell death, identifying bacterial amyloid as a central driver of non-lytic inflammasome activation during immune recognition of biofilms.

**Highlights:**

- Amyloid curli from bacterial biofilms is a novel ligand of caspase-11 and activates non canonical inflammasome without triggering pyroptotic cell death in human and mouse dendritic cells, a process called DC hyperactivation.
- Curli alone is sufficient to induce IL-1β secretion because it provides both required signals: it primes pro-IL-1β through TLR2 and TLR4 and activates the inflammasome through caspase-11.
- Curli induces a limited oligomerization of mature Gasdermin-D that localizes selectively to the plasma membrane, without targeting mitochondria or undergoing NINJ1 oligomerization.
- Curli-containing biofilms induce similar IL-1β secretion in viable DCs, highlighting DC hyperactivation as pivotal response during host recognition of bacterial biofilms.

## Introduction

Bacterial biofilms are structured microbial communities embedded in an extracellular polymeric matrix that protects bacteria from a hostile environment, confer resistance to antibiotics and host immune clearance, driving chronic and recurrent infections that are difficult to eradicate ^1–5^. Beyond direct infectious complications, persistent biofilm-associated inflammation has been implicated in the pathogenesis of autoimmunity, atherosclerosis, and cancer ^6–9^, underscoring the importance of understanding how the immune system recognizes biofilm components.

The biofilm matrix contains polysaccharides, lipids, extracellular nucleic acids, and functional amyloid fibers ^4^. Among the latter, most notable is curli, produced by *Enterobacteriaceae* including *Escherichia coli* and *Salmonella* species ^10–13^. Curli amyloids are extracellular polymeric fibrils with the characteristic cross-β-sheet structure of amyloids that are essential for the architecture and stability of *Enterobacteriaceae* biofilms ^11^. In addition to their structural role, curli function as potent pattern-associated molecular pattern (PAMP) recognized by host pattern recognition receptors (PRRs), including Toll-like receptor (TLR) 2 ^14,15^. We previously demonstrated that curli induce robust innate immune responses in murine macrophages and dendritic cells (DCs), and that curli-driven immune activation promotes autoimmunity ^6,15^. Specifically, curli-producing bacteria accelerate disease progression in murine lupus, while anti-curli antibody levels correlate with disease flares in patients with systemic lupus erythematosus, establishing a mechanistic link between biofilm-associated PAMPs and autoimmune pathology ^6,7^. Among the innate immune cells that sense curli, DCs are uniquely positioned to bridge innate and adaptive immunity by translating pathogen recognition into antigen-specific T-cell responses ^16,17^. Despite their central role in orchestrating immune responses, the molecular mechanisms by which DCs recognize and respond to biofilm-associated PAMPs such as curli remain incompletely understood.

IL-1β promotes Th17 responses ^18^, which are considered pivotal immune response against extracellular bacteria ^19,20^. IL-1β production requires an initial TLR-dependent priming step that upregulates pro-IL-1β expression, followed by a second signal, the activation of the inflammasome ^21,22^: this multimeric protein complex is assembled upon stimulation by microbial or endogenous danger signals, culminating in the activation of inflammatory caspases ^23–25^. In the canonical pathway, sensor molecules such as NLRP3 recruit the adaptor, apoptosis-associated speck-like protein containing a CARD (ASC) and procaspase-1, whose activation drives proteolytic processing of pro-IL-1β into its biologically active form and induces pyroptosis, an inflammatory lytic cell death ^23,26–28^ ^29^.

Beyond the canonical pathway, a non-canonical inflammasome pathway operates through caspase-11 in mice and caspase-4/5 in humans ^30–33^. Active caspase-11 cleaves Gasdermin D (GSDMD), whose N-terminal fragment forms plasma membrane pores that facilitate IL-1β release and frequently culminate in pyroptotic cell death accompanied by release of alarmins including HMGB1 ^34–36^. To date, only one PAMP, lipopolysaccharide (LPS) from Gram-negative bacteria ^32^, and one DAMP, oxidized phospholipids (oxPAPC) derived from dying cells ^37^, have been definitively shown to physically bind and activate the non-canonical inflammasome. Importantly, oxidized phospholipids activate caspase-11 in DCs but, unlike LPS, do so without inducing pyroptosis, generating a unique “hyperactivated” state in which IL-1β is secreted from viable cells ^37,38^. This uncoupling of cytokine secretion from lytic death is associated with enhanced CCR7-dependent migration and superior capacity of DCs to prime durable T cell responses compared to both pyroptotic and classically activated DCs ^39,40^, suggesting that the mode of inflammasome activation has fundamental consequences for the downstream adaptive immune response.

Rapsinski et al. previously demonstrated that curli activates the NLRP3 inflammasome in bone marrow-derived macrophages, inducing caspase-1/11-dependent IL-1β secretion without pyroptotic cell death ^41^. This non-lytic phenotype is consistent with a hyperactivation-like state, yet the underlying mechanisms linking curli recognition to inflammasome assembly, the role of GSDMD in mediating cytokine secretion in the absence of lytic death, and whether this biology extends to DCs remain entirely uncharacterized. Also, in vivo evidence indicates that curli induces IL-1β production in the intestinal mucosa, though the cellular source remains to be investigated ^42^.

Here we define the molecular mechanisms by which curli activates the inflammasome and induce IL-1β secretion in DCs uncoupled from pyroptosis. We demonstrate that curli hyperactivates both murine and human DCs via the non-canonical inflammasome pathway by directly binding caspase-11 (and activating caspase-5 in humans). IL-1β secretion is dependent on GSDMD, the mature NT-GSDMD fragment undergoes weak oligomerization and localizes exclusively to the plasma membrane, sparing the mitochondria, in absence of ninjurin-1 (NINJ-1) oligomerization, thus accounting for the lack of pyroptosis ^43,44^. Furthermore, these curli-hyperactivated DCs display enhanced migratory capacity *in vivo*, suggesting augmented adaptive immune responses. Finally, our findings that curli-containing biofilms similarly stimulate IL-1β secretion in DCs without cell death highlight curli as a key driver of non-lytic inflammasome activation during the immune response to bacterial biofilms.

## Results

### Curli amyloid induces IL-1β release without pyroptosis

We first tested the ability of curli amyloid purified from *Salmonella* Typhimurium biofilms^6^ to stimulate IL-1β production in murine and human DCs. In murine DCs (muDCs)^45^, curli induced robust, dose-dependent IL-1β release after 22 hours of stimulation (Figure 1A). This response was associated with inflammasome activation, as demonstrated by cleavage of caspase-1 (p20), GSDMD (p30) and mature IL-1β (p17) (Figure 1B), and formation of ASC specks (Figure 1C) which is essential for caspase-1 autocleavage and activation ^25^. To benchmark the IL-1β secretory capacity of these DCs, we primed DCs with LPS (3 hours) followed by NLRP3 inflammasome stimulus Nigericin (2.5 hours) that elicited release of very high amounts of IL-1β, robust cleavage of caspase-1 and GSDMD, and formation of ASC specks as expected (Figure 1B) ^23,46,47^. Curli and LPS similarly primed the inflammasome, inducing comparable levels of *Il1b* and *Il18* transcripts by RNA-seq analysis (Figure S1A) and equivalent pro-IL-1β (p31) protein expression by Western blot (Figure 1B). When followed by nigericin, both curli and LPS induced similarly high levels of IL-1β release, far exceeding curli alone (Figures 1A–C). Furthermore, combined treatment with curli and LPS did not enhance IL-1β release beyond curli alone (Figure S1B), indicating that curli is a strong priming stimulus that is sufficient to drive both priming and activation of the inflammasome.

**Figure 1.**
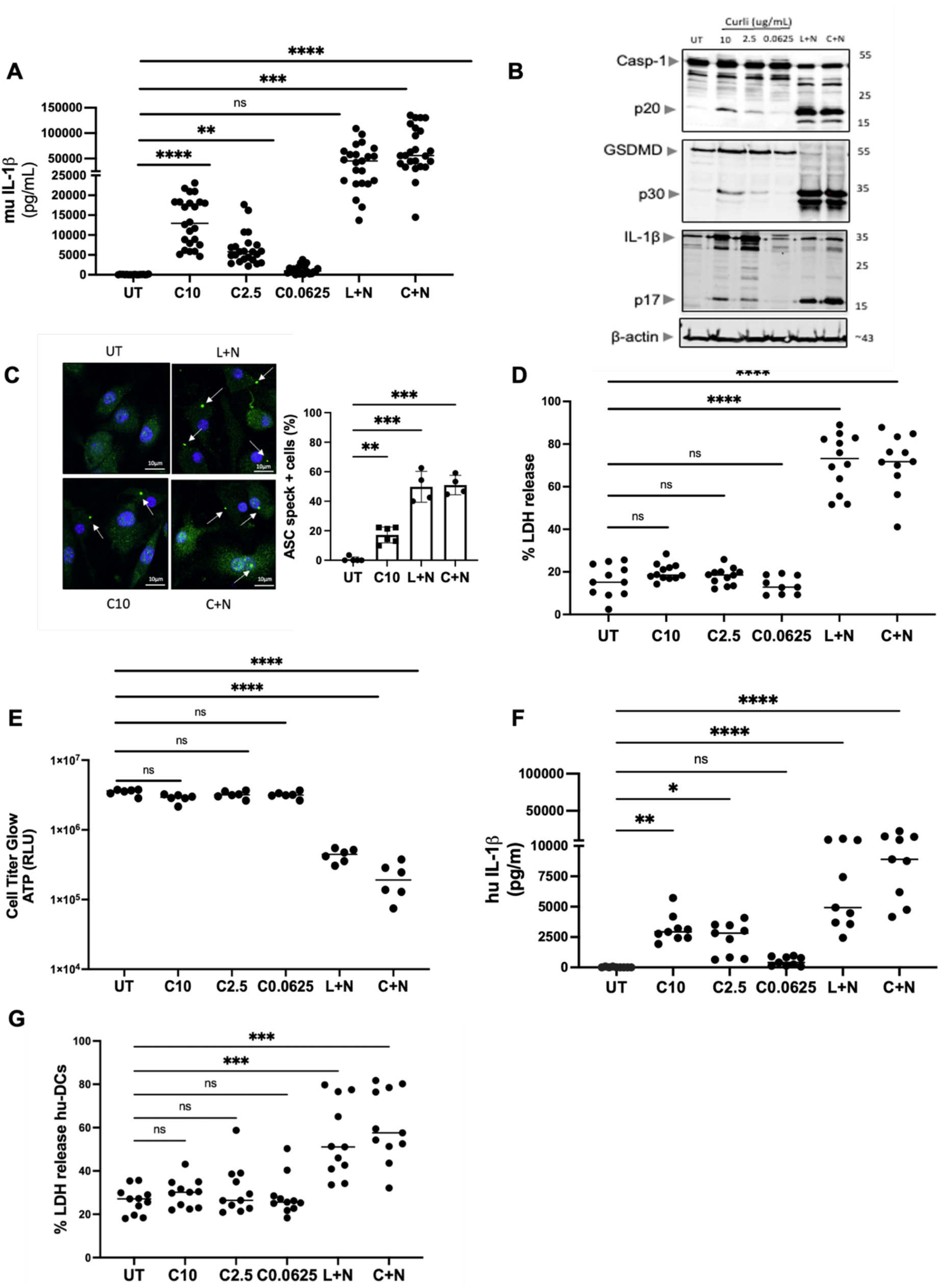
Curli activates the inflammasome without inducing cell death. (A) Murine IL-1β measured by ELISA in supernatants of WT muDCs stimulated with curli at the indicated concentrations (10, 2.5, and 0.0625 µg/mL) for 20–22 hr, or primed with LPS (100 ng/mL, 3 hr) followed by nigericin (10 µM, 2.5 hr) (L+N), or curli (10 µg/mL, 3 hr) followed by nigericin (C+N). **(B)** Immunoblot analysis of caspase-1 (pro-form and p20 cleavage fragment), GSDMD (pro-form and p30 cleavage fragment), IL-1β (pro-form and mature p17 fragment), and β-actin (loading control) in WT DCs treated with the indicated conditions. **(C)** Representative confocal immunofluorescence images (*left*) and quantification (*right*) of ASC speck formation in murine DCs under indicated conditions. ASC (green), nuclei (blue, DAPI). Arrows indicate ASC specks. Scale bars, 10 µm. Data are expressed as percentage of ASC speck-positive cells from a minimum of 150 cells per condition counted across three independent fields. **(D)** Cell death assessed by LDH in supernatants (% of maximum) and **(E)** cell viability assessed by CellTiter-Glo luminescent ATP quantification (RLU) in WT DCs under the indicated conditions. **(F)** Human IL-1β release measured by ELISA in supernatants of hu-moDCs stimulated with the indicated conditions. **(G)** Cell death assessed by LDH release (% of maximum) in supernatants of hu-moDCs stimulated with curli at indicated concentrations, L+N, or C+N. Data show mean and dots indicate biological replicates from at least three independent experiments. Statistical significance was determined by one-way ANOVA with Šídák’s or Dunnett’s multiple comparisons post-hoc test. *p < 0.05, **p < 0.01, ***p < 0.001, ****p < 0.0001; ns, not significant.

Inflammasome activation is typically coupled to pyroptosis, a lytic form of cell death resulting from gasdermin pore formation and NINJ1-mediated plasma membrane rupture ^25^. Surprisingly, curli stimulation induced no measurable cell death compared to untreated DCs, as assessed by lactate dehydrogenase (LDH) release and ATP quantification (CellTiter-Glo assay), whereas LPS+nigericin and curli+nigericin triggered rapid pyroptosis in the majority of DCs (Figures 1D–E). Furthermore, kinetic analysis of loss of plasma membrane integrity revealed that curli did not induce cell death over time, in contrast to the rapid burst induced by LPS + nigericin (Figure S1C). Together, these findings indicate that curli activates the inflammasome through a distinct mechanism in which caspase-1 and GSDMD activations are sufficient for IL-1β processing and release but remain below the threshold required to trigger pyroptotic cell death.

To assess the human relevance of these findings, we stimulated human monocyte-derived DCs (hu-moDCs) differentiated with GM-CSF and IL-4, an established *in vitro* model of inflammatory DCs and the human counterpart of murine GM-CSF-DCs ^48^. Curli induced significant IL-1β release with minimal cell death in hu-moDCs (Figures 1F–G), mirroring the pattern observed in murine DCs. Consistent with our previous report ^6^, curli also drove robust production of multiple pro-inflammatory cytokines, and we show IL-6 as representative, in both mouse and hu-moDCs (Figures S1D–E), confirming that curli elicits full DC activation in both species. Collectively, these findings demonstrate that curli engages the inflammasome machinery to drive IL-1β processing and release in both murine and human DCs, without triggering pyroptotic cell death.

### Curli primes the inflammasome through TLR2 and TLR4

To study the priming phase of curli-driven inflammasome activation, we assessed NF-κB pathway activation and found that curli induced robust IκBα degradation, phosphorylation of IκBα and p65, and subsequent pro-IL-1β upregulation with concordant kinetics (Figure 2A), collectively indicating effective inflammasome priming ^49^. To dissect the contribution of TLR adaptor signaling to this priming phase, we generated DCs from mice deficient in the key TLR adaptor proteins MYD88 or TRIF^50^. Curli-induced inflammasome activation and IL-1β secretion were significantly reduced in both *Myd88*-KO and *Trif*-KO DCs relative to wild-type, with pro-IL-1β accumulation more dependent on *Myd88*, and GSDMD cleavage more dependent on *Trif* (Figures 2B–C). The residual IL-1β secretion observed in *Trif*-KO DCs stimulated with curli+nigericin likely reflects incomplete impairment of priming, together with the ability of nigericin to bypass TRIF-dependent licensing of GSDMD activation (Figures 2B–C). Curli-induced IL-6 secretion was more dependent on MYD88 than TRIF, whereas LDH release was unchanged (Figures S2A–B). Collectively, these results establish that MYD88 and TRIF make distinct contributions to curli-induced inflammasome activation, with MYD88 driving pro-IL-1β transcription and TRIF licensing GSDMD for activation.

**Figure 2.**
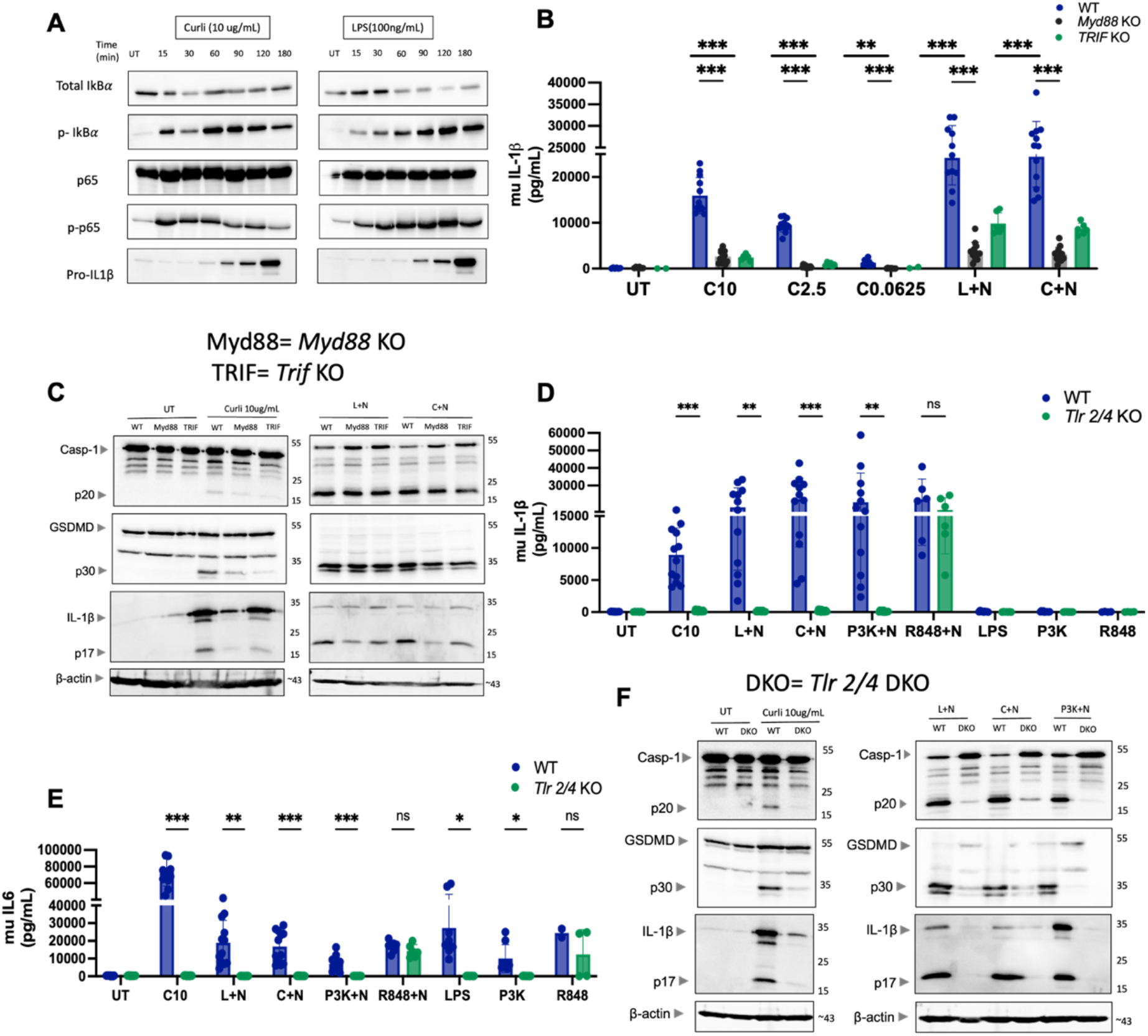
Curli primes the inflammasome through MYD88 and TRIF triggered by TLR2 and TLR4. **(A)** Immunoblot time-course analysis of NF-κB pathway activation in WT muDCs stimulated with curli (10 µg/mL) or LPS (100 ng/mL) for the indicated times (0–180 min). Blots show total IκBα, phospho-IκBα, total p65, phospho-p65, and pro-IL-1β. **(B)** Murine IL-1β measured by ELISA in WT (blue), *Myd88* KO (black), and *Trif* KO (green bars) muDCs stimulated with curli at the indicated concentrations for 20–22 hr, L+N, or C+N for 3+2.5 hrs. **(C)** Immunoblot analysis of caspase-1 (pro-form and p20 cleavage fragment), GSDMD (pro-form and p30 cleavage fragment), IL-1β (pro-form and mature p17 fragment), and β-actin (loading control) in WT, *Myd88* KO, and *Trif* KO BMDCs under the indicated conditions. **(D)** Murine IL-1β in WT (blue) and *Tlr2/4* DKO (green) muDCs stimulated with curli (10 µg/mL), L+N, C+N, Pam3CSK4 (500ng/ml) + nigericin (P3K+N), R848 5ug/ml + nigericin (R848+N), LPS, Pam3CSK4 (P3K), or R848 alone. **(E)** Murine IL-6 measured by ELISA in WT (blue) and *Tlr2/4* DKO (green) muDCs under the indicated conditions. **(F)** Immunoblot analysis of caspase-1, GSDMD, IL-1β, and β-actin in WT and *Tlr2/4* DKO muDCs stimulated with curli (10 µg/mL), L+N, C+N, or P3K+N. Data represent mean ± SEM from at least three independent experiments, dots indicate biological replicates. Statistical significance was determined by multiple unpaired t-tests with Šídák’s correction for multiple comparisons between WT and KO genotypes within each treatment condition. *p < 0.05, **p < 0.01, ***p < 0.001, ****p < 0.0001; ns, not significant.

The combined deletion of TLR2 and TLR4 was required to abolish curli-induced IL-1β and IL-6 production (Figures 2D–E), as well as the IL-1β, IL-6, and cell death responses elicited by curli+nigericin and by canonical TLR agonists (LPS for TLR4; Pam3CSK4 for TLR2) paired with nigericin (Figure S2C). By contrast, responses to the TLR7 ligand R848+nigericin were preserved in *Tlr2/4*-double KO (DKO) DCs, confirming the specificity of TLR2/TLR4 engagement by curli (Figures 2D–E and Figure S2C). Analysis of single-knockout DCs demonstrated distinct and partially overlapping contributions of TLR2 and TLR4 to pro-IL-1β priming (Figures S2D–G). Consistent with this, upregulation and cleavage of pro-IL-1β were markedly reduced in *Tlr2/4*-double KO DCs, and caspase-1 and GSDMD cleavage was also impaired, while their expression was not impacted, reflecting the fact that both proteins are basally expressed in GM-CSF-differentiated DCs (Figure 2F). Concordant results were obtained in *Tlr2*-KO DCs treated with the selective TLR4 inhibitor TAK-242^51^, pharmacologically recapitulating the double-knockout phenotype (Figures S2D–F).

Since curli was purified from *Salmonella* Typhimurium biofilms, preparations inevitably contain some amounts of LPS. To minimize potential LPS confounding, we utilized the *msbB* mutant of *Salmonella* Typhimurium, which produces a penta-acylated Lipid A that fails to engage human TLR4 and is a weak agonist of murine TLR4 ^52–54^. To directly assess whether curli, rather than contaminating LPS, is responsible for TLR4 activation, we stimulated murine DCs with curli in the presence of polymyxin B, a cyclic peptide antibiotic that sequesters LPS and prevents its binding to TLR4^55,56^. Curli induced equivalent levels of IL-1β and IL-6 regardless of polymyxin B co-treatment, demonstrating that TLR4 activation by curli occurs independently of LPS contamination (Figures S3A-B). Taken together, these findings establish that TLR2 and TLR4 act cooperatively through MYD88 and TRIF to drive curli-induced inflammasome priming and downstream cytokine production via NF-κB.

### Curli induces IL-1β release via NLRP3 inflammasome

To characterize the molecular mechanisms underlying curli-induced inflammasome activation, we assessed IL-1β secretion and inflammasome component processing in *Nlrp3*-, *Asc*-, and *Casp1*-KO DCs, and in WT DCs treated with the selective NLRP3 inhibitor MCC950^57^. Curli induced robust IL-1β release in WT DCs, whereas all three KOs and MCC950-treated WT DCs exhibited strongly reduced or absent IL-1β secretion and no detectable cell death (Figures 3A, S4A–B), establishing that curli-driven IL-1β release depends on canonical NLRP3 inflammasome assembly. Consistent with a role for potassium efflux as the upstream activation signal, supplementation of extracellular potassium dose-dependently suppressed curli-induced IL-1β release without affecting cell viability or DC function (Figures 3C, S4C–D), placing curli within the canonical class of NLRP3 stimuli that operate through K⁺ efflux ^22,23,46,47,58^.

**Figure 3.**
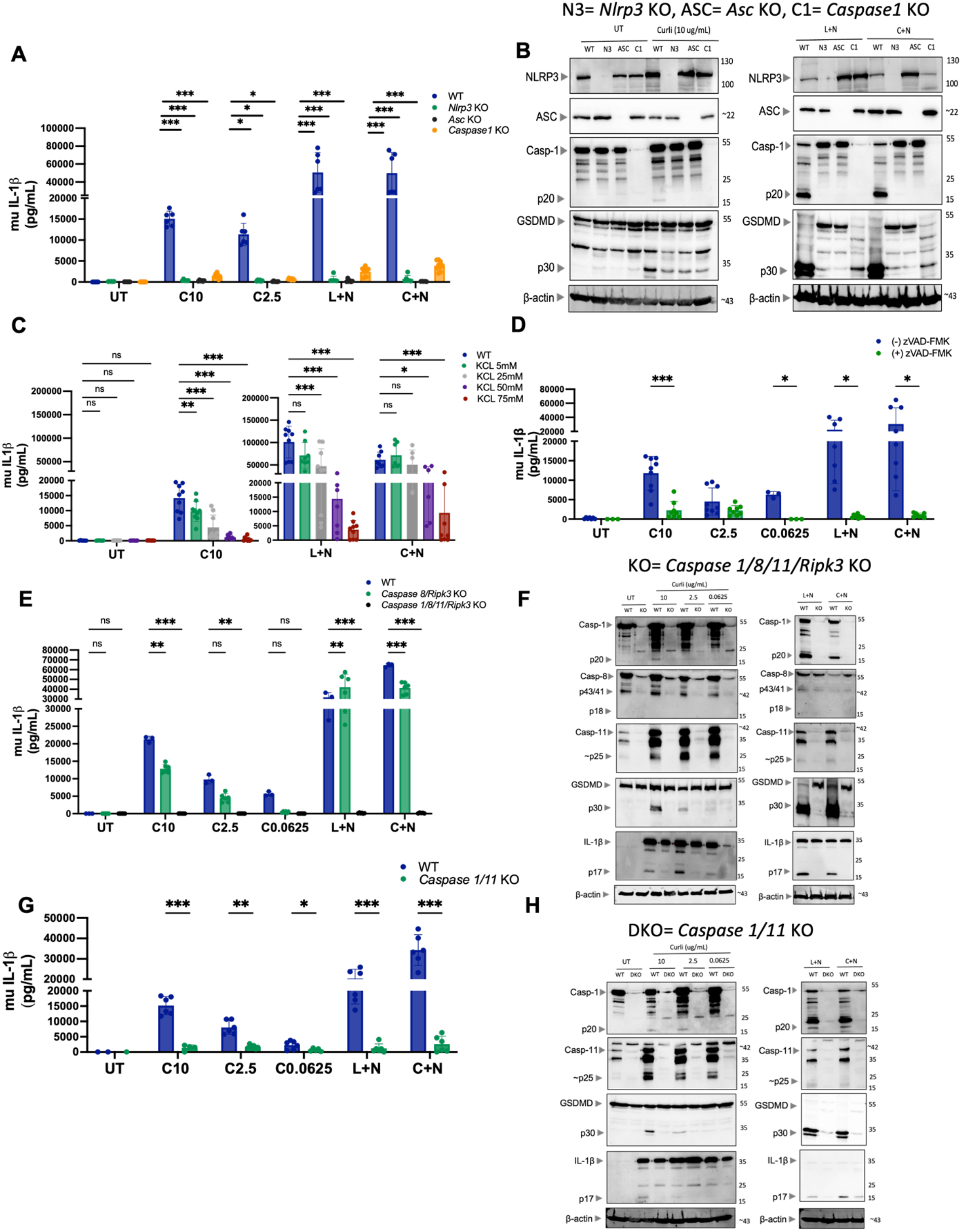
Curli-induced inflammasome activation is dependent on caspases 1/11 and NLRP3. **(A)** Murine IL-1β measured by ELISA in supernatants of WT (blue), *Nlrp3* KO (green), *Asc* KO (black), and *Caspase1* KO (orange) muDCs stimulated with curli (10, 2.5 µg/mL), L+N, or C+N. **(B)** Immunoblot analysis of NLRP3, ASC, caspase-1, GSDMD and β-actin in WT, *Nlrp3* KO (N3), *Asc* KO (ASC), and *Caspase1* KO (C1) muDCs stimulated with curli (10 µg/mL), L+N, or C+N. **(C)** Murine IL-1β in supernatants of WT muDCs pre-treated with increasing concentrations of extracellular KCl (5, 25, 50, 75 mM) prior to stimulation with curli (10 µg/mL), L+N, or C+N, demonstrating dose-dependent suppression of curli-driven inflammasome activation by extracellular potassium. **(D)** Murine IL-1β in supernatants of WT muDCs stimulated with curli (10, 2.5, 0.0625 µg/mL), L+N, or C+N in the presence (+) or absence (−) of the pan-caspase inhibitor zVAD-FMK (20 µM). **(E)** Murine IL-1β in supernatants of WT (blue), *Caspase 8/Ripk3* DKO (teal), and *Caspase 1/8/11/Ripk3* KO (green) muDCs stimulated with curli at the indicated concentrations, L+N, or C+N. **(F)** Immunoblot analysis of caspase-1 (pro-form and p20), caspase-8 (pro-form, p43/41, and p18), caspase-11, GSDMD, IL-1β and β-actin in WT and *Caspase 1/8/11/Ripk3* KO muDCs stimulated with curli, L+N, or C+N. **(G)** Murine IL-1β in supernatants of WT (blue) and *Caspase 1/11* DKO (green) muDCs stimulated with curli, L+N, or C+N. **(H)** Immunoblot analysis of caspase-1, caspase-11, GSDMD, IL-1β and β-actin in WT and *Caspase 1/11* DKO muDCs stimulated with curli, L+N, or C+N. Data represent mean ± SEM from at least three independent experiments, dots indicate biological replicates. Statistical significance was determined by multiple unpaired t-tests with Šídák’s correction or two-way ANOVA where appropriate. *p < 0.05, **p < 0.01, ***p < 0.001, ****p < 0.0001; ns, not significant.

Notably, while caspase-1 cleavage was absent in all KO DCs, confirming engagement of the canonical NLRP3 pathway, GSDMD cleavage was still observed (Figure 3B), implicating the involvement of additional caspases in GSDMD processing upon curli stimulation^59^. Consistent with this, the pan-caspase inhibitor zVAD-FMK significantly reduced curli-induced mature IL-1β secretion (Figure 3D).

To identify the caspase(s) responsible for caspase-1-independent GSDMD cleavage, we utilized DCs from multiple KO mice lacking caspases known to process GSDMD: a quadruple KO of caspase-1, -8, and -11 with RIPK3; a caspase-8/RIPK3 double KO (DKO) ^60–63^, and a caspase-1/11 DKO. Curli-induced IL-1β secretion and inflammasome activation were dependent on both caspase-1 and caspase-11 but were fully independent of caspase-8 (Figures 3E–H, S4E– I). Accordingly, robust caspase-11 cleavage was detected in WT DCs stimulated with curli, whereas caspase-8 cleavage was absent, and GSDMD processing into its active N-terminal fragment was specifically lost in genotypes lacking caspase-11 but not in caspase-8/RIPK3 DKO DCs (Figures 3F, 3H, S4I). IL-6 production was induced by curli unaffected across all KO DCs, confirming its independence from inflammasome signaling (Figures S4F–G).

Collectively, these data establish that curli activates the canonical NLRP3 inflammasome via K⁺ efflux and drives IL-1β secretion through a mechanism specifically requiring both caspase-1 and caspase-11, without involvement of caspase-8.

### Curli directly binds caspase-11 and induces IL-1β release via non-canonical inflammasome activation

To directly investigate the role of caspase-11 in curli-induced IL-1β production, we compared WT and *Casp11*-KO DCs and observed a strong reduction in curli-induced IL-1β secretion, accompanied by time-dependent attenuation of caspase-1, GSDMD, and mature IL-1β cleavage in *Casp11*-KO DCs (Figures 4A–B), without changes in IL-6 secretion or cell death (Figures S5A–B). Transfected LPS, a well-established activator of the non-canonical inflammasome via cytosolic caspase-11 ^30–33^ was used as a positive control. Consistent with impaired upstream caspase-11 activity, curli-induced ASC speck formation was also significantly reduced in *Casp11*-KO DCs (Figure 4C).

**Figure 4.**
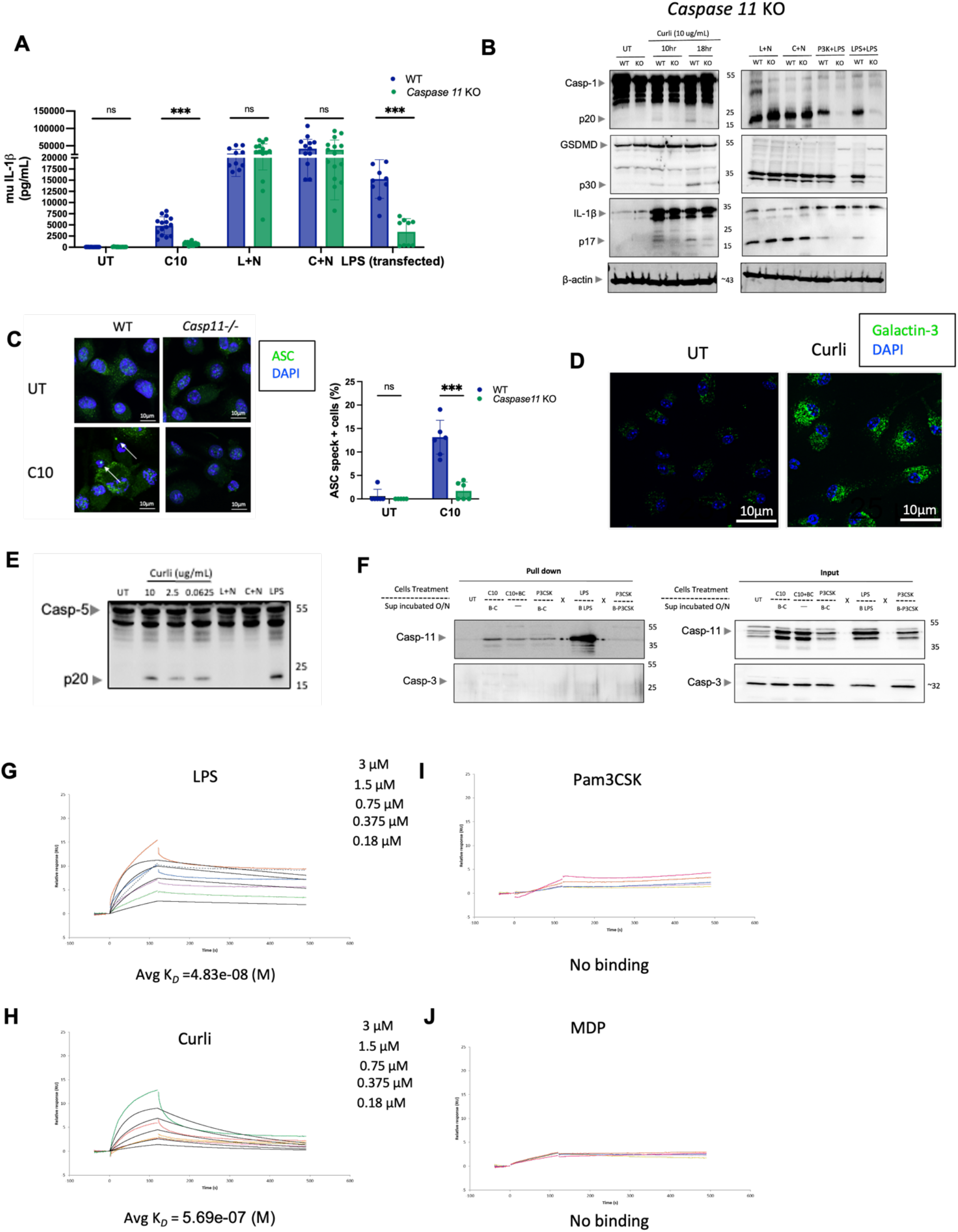
Curli directly binds caspase-11 and activates the non-canonical inflammasome. **(A)** Murine IL-1β measured by ELISA in supernatants of WT (blue) and *Caspase 11* KO (green) muDCs stimulated with curli (10 µg/mL), L+N, C+N, or transfected LPS (positive control for non-canonical caspase-11 activation; 6hrs). **(B)** Immunoblot analysis of caspase-1, GSDMD, IL-1β, and β-actin in WT and *Caspase 11* KO muDCs stimulated with curli (10 µg/mL) at 10 and 18 hr, L+N, C+N, P3K+LPS, or transfected LPS. **(C)** Representative confocal immunofluorescence images (*left*) and quantification (*right*) of ASC speck formation in WT and *Casp11*^⁻/⁻^ muDCs stimulated with curli (10 µg/mL) or left untreated. ASC (green), nuclei (blue, DAPI). Arrows indicate ASC specks. Scale bars, 10 and 30 µm. Data in bar graph are expressed as percentage ASC speck-positive cells from a minimum of 150 cells per condition. **(D)** Representative confocal immunofluorescence images of Galectin-3 (green) and nuclei (blue, DAPI) in muDCs under the indicated conditions. Scale bars, 10 µm. **(E)** Immunoblot analysis of caspase-5 (pro-form and p20 cleavage fragment) in hu-moDCs stimulated with curli (10, 2.5, 0.0625 µg/mL), L+N, C+N, or LPS. **(F)** muDCs were primed with LPS, Pam3CSK4, curli (3 hr), or curli plus biotinylated curli. Lysates were then incubated overnight with the corresponding biotinylated ligand (biotinylated LPS, positive control; biotinylated Pam3CSK4, negative control; or biotinylated curli) plus streptavidin beads, except for the curli-plus-biotinylated-curli condition, which received streptavidin beads alone. Pull-down (left) and input (right) blots were probed for caspase-11 and caspase-3, demonstrating specific binding of curli and LPS to caspase-11 but not caspase-3. X denotes empty spacer wells. **(G–J)** Surface plasmon resonance (SPR) showing direct binding of recombinant caspase-11 immobilized on sensor chip to LPS (avg KD∼ = 4.83 × 10⁻⁸ M) **(G)**, curli (avg KD∼ = 5.69 × 10⁻⁷ M) **(H)**, Pam3CSK4 (no binding detected) **(I)**, and MDP (no binding detected) **(J)**, at analyte concentrations of 0.18–3 µM. Data represent mean ± SEM from at least three independent experiments, dots indicate biological replicates. Statistical significance determined by unpaired t-tests (Mann-Whitney) or two-way ANOVA. *p < 0.05, **p < 0.01, ***p < 0.001, ****p < 0.0001; ns, not significant.

Since caspase-11 is localized in the cytosol, we investigated whether curli could access this compartment following internalization. Following 6–8 hours of incubation, Congo Red-stained curli was detected within intracellular vesicles, a subset of which co-localized with the lysosomal marker LAMP-2, while fewer vesicles were positive for the early endosomal marker Rab5, suggesting that curli undergoes endocytosis within the first few hours of exposure and traffics to the lysosomes (Figures S5C–D). Immunofluorescence analysis of Galectin-3 puncta, a marker of lysosomal membrane damage, revealed a marked increase in puncta formation upon curli stimulation compared to untreated controls (Figure 4D), suggesting that curli disrupts the phago-lysosomal membrane to gain access to the cytoplasm and engage caspase-11. Supporting translational relevance, we observed dose-dependent processing and activation of caspase-5, the human orthologue of murine caspase-11, in hu-moDCs stimulated with increasing concentrations of curli, mirroring the response to transfected LPS and indicating conservation of non-canonical inflammasome engagement across species (Figure 4E).

To determine whether curli directly interacts with caspase-11, we performed co-immunoprecipitation (Co-IP) using biotinylated curli, with biotinylated LPS and Pam3CSK4 as positive and negative controls, respectively ^32^. Similar to LPS, curli binds caspase-11 but not caspase-3, confirming specificity (Figure 4F). To quantify this interaction, we performed surface plasmon resonance (SPR) and measured direct binding of curli to caspase-11 with a mean Kᴅ ≈ 5.69 × 10⁻⁷ M, comparable to LPS (Avg Kᴅ ≈ 4.83× 10⁻⁸) (Figures 4G, H). In contrast, the TLR2 ligand Pam3CSK4 and the NOD2 ligand muramyl dipeptide (MDP) showed no detectable binding to caspase-11 as expected, confirming the selectivity of this interaction (Figures 4I, J).

Overall, these findings demonstrate that curli directly binds caspase-11 with strong affinity, engaging the non-canonical inflammasome pathway in both murine and human DCs. In murine DCs, curli activated caspase-11 induces GSDMD cleavage, triggering potassium efflux that activates the NLRP3-caspase-1 axis and drives IL-1β release.

### Recombinant CsgA fibers as a defined tool to study non-canonical caspase-11-dependent inflammasome activation

To confirm that caspase-11 binding and inflammasome activation are intrinsic properties of curli rather than a consequence of co-purifying contaminants, we synthesized recombinant full length CsgA, the major monomeric subunit of *Salmonella* curli fibers ^6,11^, and polymerized it into amyloid fibrils *in vitro*. We confirmed the formation of fibers by Thioflavin T staining (ThT) assay and analyzed fiber morphology and size parameters, known to influence the potency of innate immune stimulation^64,65^ (Figures S6A-D): CsgA fibers showed similar features to natural curli.

Stimulation of WT muDCs with CsgA fibers recapitulated the responses observed with natural curli: robust IL-1β secretion (1–5 ng/ml) was induced without detectable cell death, which was not further altered by a five-fold increase in CsgA fiber dose (Figures 5A–C). CsgA fibers also up-regulated the expression of caspase-11 and pro-IL-1β and triggered cleavage of caspase-11, caspase-1, and GSDMD, and drove mature IL-1β processing, collectively mirroring the non-canonical inflammasome activation signature of natural curli (Figure 5B). Co-immunoprecipitation showed that CsgA fibers interacted with caspase-11 but not caspase-3, reproducing the binding specificity observed with natural curli (Figure 5D). SPR analysis further demonstrated that CsgA fibers binds caspase-11 directly with high affinity (KD = 6.045 ×10⁻⁷ M) comparable to that measured for natural curli (Figure 5E).

**Figure 5.**
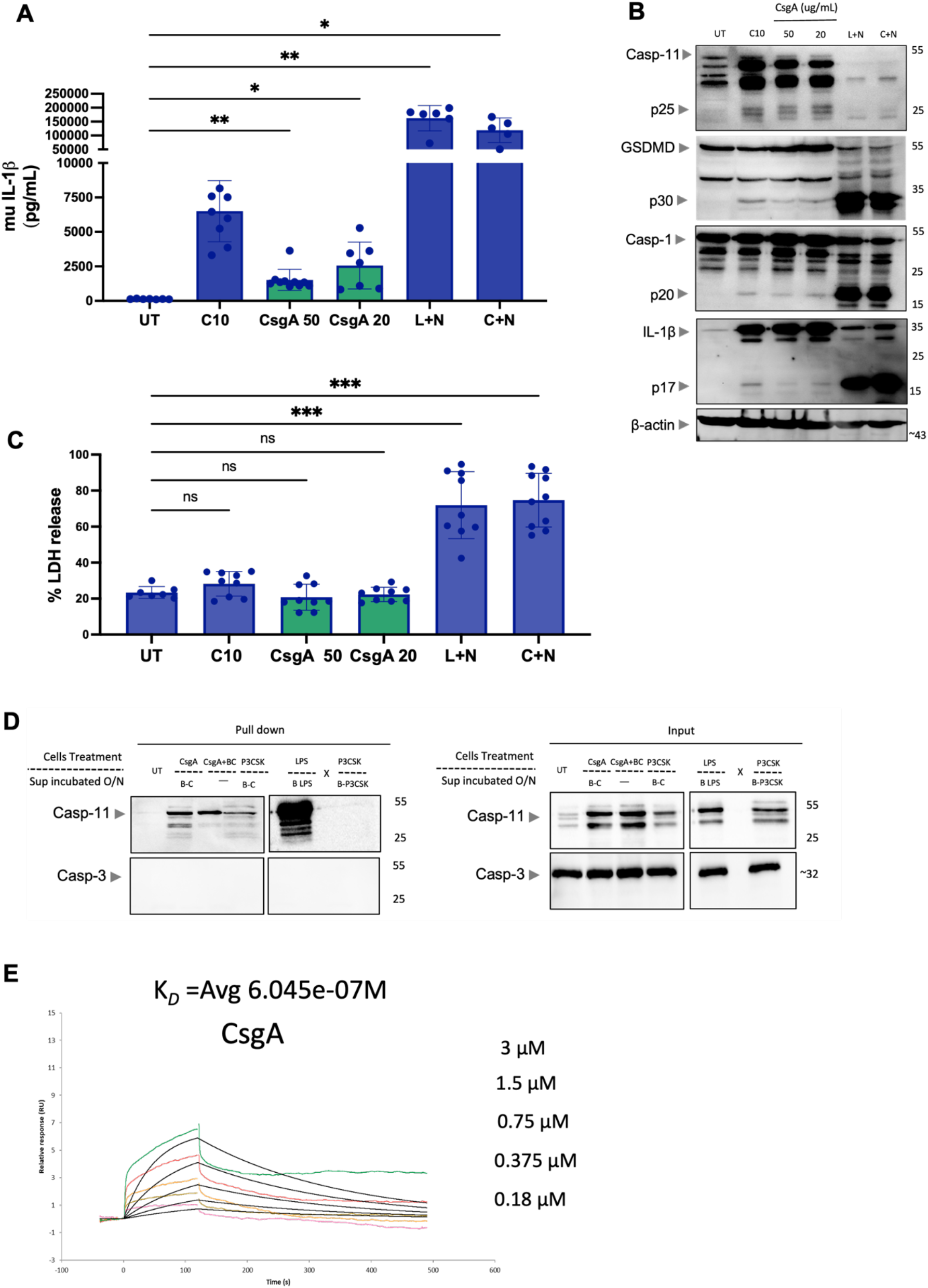
CsgA fibers, fibrillized in vitro from the monomeric subunit of curli, activate the inflammasome in a similar manner as curli. **(A)** Murine IL-1β in supernatants, **(B)** Immunoblot analysis of caspase-11, GSDMD, caspase-1, IL-1β and β-actin in lysates, and **(C)** Cell death (% LDH release) of WT muDCs stimulated with natural curli (10 µg/mL), CsgA fibers (50 or 20 µg/mL) for 2-22 hrs, and L+N, or C+N for 3+2.5 hours. **(D)** Co-immunoprecipitation pull-down using biotinylated CsgA fibers (with CsgA priming), biotinylated LPS (positive control), or biotinylated Pam3CSK4 (negative control) incubated overnight. Pull-down (*left*) and input (*right*) blots probed for caspase-11 and caspase-3, confirming specific binding of CsgA fibers to caspase-11 but not caspase-3. X denotes empty spacer wells. **(E)** SPR showing direct binding of recombinant CsgA fibers to immobilized caspase-11 at concentrations of 0.18–3 µM (avg K∼D∼= 6.045 × 10⁻⁷ M), comparable to the binding affinity of natural curli. Data represent mean ± SEM from at least three independent experiments, dots indicate biological replicates. Statistical significance determined by one-way ANOVA. *p < 0.05, **p < 0.01, ***p < 0.001, ****p < 0.0001; ns, not significant.

Collectively, these results demonstrate that recombinant CsgA fibers fully phenocopies the inflammasome-activating activity of naturally purified curli, establishing that the response is driven by the curli amyloid fiber itself rather than co-purifying biofilm contaminants.

### Curli induces Gasdermin D-dependent IL-1β release in absence of pyroptosis

Curli-induced GSDMD cleavage, detected in Figures 1B, 3B, 4B, and 5B, prompted us to directly assess the requirement of GSDMD for curli-driven inflammasome activation and IL-1β release, which may be mediated by other gasdermins expressed by DCs, such as Gasdermin E. IL-1β release was significantly reduced in *Gsdmd*-KO compared to WT DCs stimulated with curli, while no cell death was observed in either genotype (Figures 6A, S7A). Interestingly, caspase-1 cleavage occurred exclusively in WT DCs, indicating that caspase-1 processing is downstream of and dependent on GSDMD, whereas cleaved caspase-11 was detected in both WT and *Gsdmd*-KO DCs, placing caspase-11 activation upstream of and independent of GSDMD (Figure 6B). The robust IL-1β release and cell death elicited by curli+nigericin, comparable in magnitude to LPS+nigericin-were both significantly reduced in *Gsdmd*-KO DCs (Figures 6A, S6A), further underscoring the central role of GSDMD in curli-induced inflammasome outputs.

**Figure 6.**
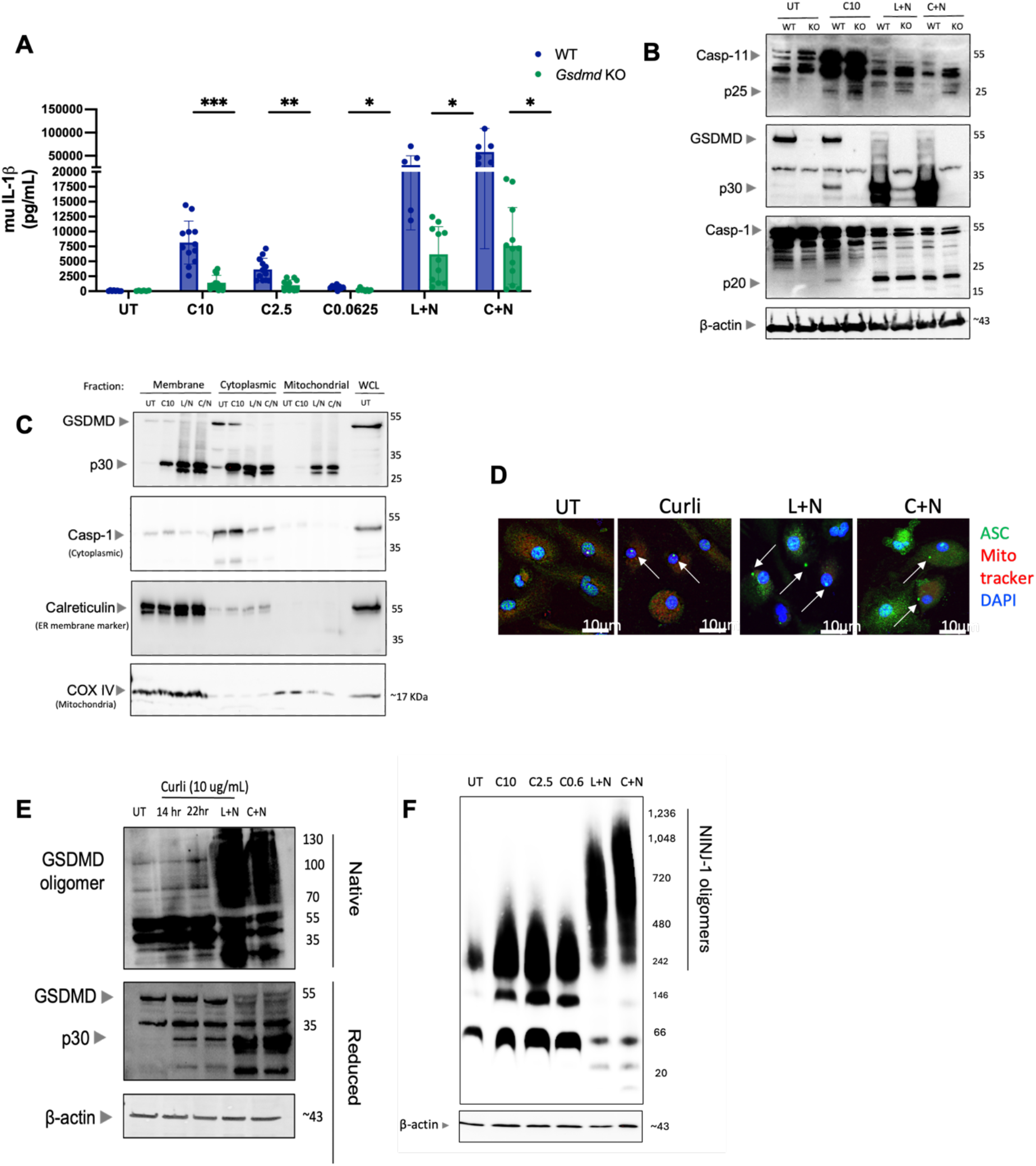
Curli-mediated IL-1β secretion is GSDMD-dependent. **(A)** Murine IL-1β measured by ELISA in supernatants and **(B)** Immunoblot analysis of caspase-11, GSDMD, caspase-1 and β-actin in lysates of WT and *Gsdmd* KO muDCs stimulated with curli (C10), L+N, or C+N. **(C)** Subcellular fractionation immunoblot showing distribution of GSDMD (pro-form and p30) across membrane, cytoplasmic, and mitochondrial fractions in WT muDCs stimulated with curli (C10), L+N, or C+N. Caspase-1 (cytoplasmic fraction marker), calreticulin (ER/membrane marker), and COX IV (mitochondrial marker) confirm fraction purity. Whole cell lysate (WCL) from untreated cells shown as reference. **(D)** Representative confocal immunofluorescence images of muDCs left untreated or stimulated with curli (10 µg/mL), L+N, or C+N, co-stained for ASC (green), Mito Tracker (red), and nuclei (blue, DAPI). Arrows indicate ASC specks. Loss of Mito Tracker signal in L+N and C+N conditions indicates mitochondrial depolarization. Scale bars, 10 µm. **(E)** Native PAGE (upper) and reducing SDS-PAGE (lower) immunoblot analysis of GSDMD oligomerization in WT muDCs stimulated with curli (10 µg/mL) at 14 and 22 hr, L+N, or C+N at 3+2.5 hrs. Native PAGE detects high-molecular-weight GSDMD oligomers indicative of pore assembly; reducing conditions confirm GSDMD-NT (p30) cleavage. β-actin shown as loading control. **(F)** Blue native PAGE analysis of NINJ1 oligomeric complexes in WT muDCs stimulated with curli, L+N, or C+N. β-actin shown as loading control. Data represent mean ± SEM from at least three independent experiments, dots indicate biological replicates. Statistical significance determined by unpaired t-tests with Šídák’s correction between WT and *Gsdmd* KO within each stimulation group. *p < 0.05, **p < 0.01, ***p < 0.001, ****p < 0.0001; ns, not significant.

We next investigated why GSDMD cleavage upon curli stimulation does not culminate in pyroptosis. Previous studies have established that inflammasome-induced pyroptosis requires translocation of the GSDMD N-terminal (p30) fragment to mitochondria, resulting in loss of mitochondrial membrane potential ^44^, and to the plasma membrane to form pores. Subcellular fractionation revealed that upon curli stimulation, the GSDMD-p30 fragment, basally present in the cytoplasmic fraction, partially redistributed to membrane fractions containing the plasma membrane, but was virtually absent from the mitochondrial fraction. In contrast, mitochondrial GSDMD accumulation was clearly detected in DCs undergoing pyroptosis following LPS+nigericin or curli+nigericin treatment (Figure 6C). Complementary Mito Tracker staining confirmed that curli-stimulated DCs formed ASC specks, indicative of active inflammasome assembly, while retaining mitochondrial membrane potential and morphological integrity, consistent with functional mitochondria and cell viability. LPS+nigericin and curli+nigericin, by contrast, abolished Mito Tracker signal, consistent with imminent pyroptotic cell death (Figure 6D). These observations suggest that curli drives limited GSDMD pore formation at the plasma membrane that is sufficient to permit cytokine release but falls below the threshold required to trigger cell death^44^. Supporting this interpretation, blue native PAGE analysis detected only weak GSDMD oligomerization after curli stimulation, reaching the sensitivity threshold of the assay only at 22 hours, and was markedly attenuated compared to the robust oligomers induced by LPS+nigericin and curli+nigericin. Immunoblotting under reducing conditions confirmed the presence of cleaved GSDMD-NT, indicating that limited but real pore-forming activity is engaged by curli (Figure 6E)^35^.

Upon pyroptosis triggering, the plasma membrane protein ninjurin-1 (NINJ1) forms large oligomers required for terminal cell rupture following inflammasome activation ^43,66,67^. Blue native PAGE analysis revealed that curli stimulation induced only small NINJ1 oligomeric complexes (< 480 kDa), in stark contrast to the high-molecular-weight complexes (> 480 kDa, up to 1,236 kDa) assembled in response to LPS+nigericin and curli+nigericin (Figure 6F), providing a molecular confirmation that pyroptosis is not occurring under curli stimulation conditions.

Collectively, these data demonstrate that GSDMD cleavage and limited oligomerization are essential for curli-induced IL-1β secretion from viable DCs, defining a non-pyroptotic mode of non-canonical inflammasome activation. The absence of GSDMD translocation to mitochondria, the preservation of mitochondrial membrane potential, and the failure to assemble large NINJ1 oligomeric complexes together provide a mechanistic framework describing how curli-stimulated DCs sustain inflammasome activation and cytokine release while avoiding pyroptotic cell death.

### Curli-containing bacterial biofilms hyperactivate DCs

Curli is a crucial structural component of *Enterobacteriaceae* biofilms^10,68^. To determine whether curli hyperactivates DCs in the context of a biofilm, we generated biofilms using *msbB* mutant ^52,53^ of Uropathogenic *E. coli* UTI89 strain, which expresses curli fibers (Figures S8A), and then exposed murine and human DCs to these biofilms and measured IL-1β secretion. To prevent direct bacterial infection of DCs and depletion of culture nutrients, biofilms were pre-treated with gentamicin to halt bacterial growth. Biofilm elicited robust IL-1β secretion in both cell types, demonstrating that curli-containing biofilm engages the inflammasome pathway (Figures 7A–B). No significant cytotoxicity was observed with the biofilm in either murine or human DCs (Figures 7C–D), and IL-6 secretion was comparable to that induced by purified curli alone (Figures S8B–C). Together, these results indicate that the inflammasome-activating capacity of curli occurs within the biofilm context and is neither suppressed nor antagonized by other biofilm components.

**Figure 7.**
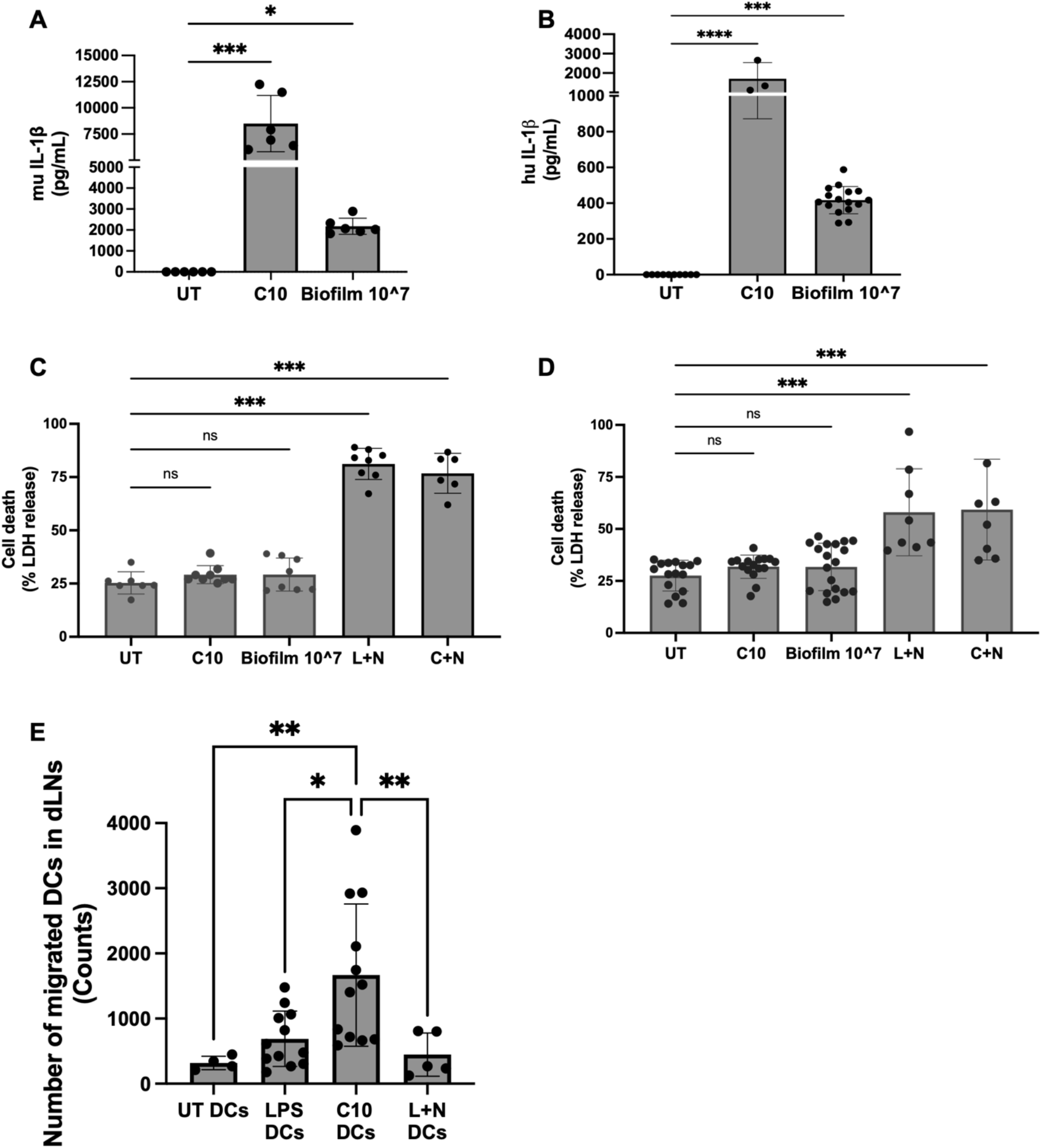
Bacterial biofilms hyperactivate DCs comparably to curli, and curli-hyperactivated DCs display enhanced migration to the draining lymph nodes. (**A)** Murine IL-1β release by ELISA in WT muDCs stimulated with curli (10 µg/mL) or UPEC *msbB* biofilm (10^7 eCFU), gentamicin-treated to stop bacteria growth prior to DC stimulation. **(B)** Human IL-1β release by ELISA in hu-moDCs stimulated with curli (10 µg/mL) or UPEC *msbB* biofilm. **(C)** Cell death (% LDH release) in WT muDCs stimulated with curli (10 µg/mL), UPEC biofilm, L+N, or C+N. **(D)** Cell death (% LDH release) in hu-moDCs stimulated with curli (10 µg/mL), biofilm, L+N, or C+N. **(E)** Quantification of CTV-labeled BMDCs recovered from draining popliteal lymph nodes 36 hr after subcutaneous footpad injection into syngeneic C57BL/6 mice. muDCs were pre-stimulated *in vitro* with curli (10 µg/mL), LPS (100 ng/mL), or L+N prior to injection; unstimulated CTV-labeled DCs served as negative control. Data presented as counts of CTV⁺ donor DCs recovered per lymph node. Data represent mean ± SEM from at least three independent experiments, dots indicate biological replicates, with five independent experiments for (**E**). Statistical significance in **E** was determined by one-way ANOVA with Tukey’s post-hoc test. *p < 0.05, **p < 0.01, ***p < 0.001, ****p < 0.0001; ns, not significant. See also Figure S7.

### Curli-hyperactivated DCs exhibit enhanced migration to the draining lymph nodes

To assess the functional consequences of curli-induced DC hyperactivation, we examined DC migration to draining lymph nodes *in vivo*, one of the earliest and most critical DC functions following PAMP recognition^69^. Cell Trace Violet (CTV)-labeled murine DCs were stimulated *in vitro* and injected into the footpad of syngeneic mice, and we quantified them by Flow Cytometry in the draining popliteal lymph nodes 24–48 hours post-injection. Curli-stimulated DCs exhibited markedly enhanced migration compared to both untreated and LPS-stimulated DCs, with an average 5-fold and 2.4-fold increase in lymph node-resident DC numbers, respectively (Figure 7E). By contrast, DCs stimulated with the pyroptotic stimulus LPS+nigericin migrated at levels comparable to untreated DCs, likely reflecting the minority of activated DCs that escaped pyroptosis (Figure 7E). Collectively, these results demonstrate that the non-pyroptotic mode of inflammasome activation induced by curli confers a functional advantage to DCs by promoting their migration to draining lymph nodes, a response that is not recapitulated by stimuli that couple inflammasome activation to pyroptotic cell death.

## Discussion

Our study demonstrates that curli, an amyloid major structural component of bacterial biofilms produced by enteric pathogens like *E. coli* and *Salmonella* ^10,70^, is a potent inflammasome stimulus that drives robust IL-1β secretion in viable DCs, a phenotype consistent with DC hyperactivation ^37,38^. Importantly, we show that full biofilms recapitulate this response, arguing that DC hyperactivation is engaged during biofilm encounter and not merely in the presence of purified amyloid. While DC hyperactivation was initially described as a feature of sterile inflammation induced by endogenous danger signals such as OxPAPC and PGPC ^37^, our results demonstrate that it can be triggered by a bacterial PAMP, suggesting it as a physiologically relevant response in chronic and recurrent biofilm-associated infections. Our results open a new perspective of the recognition of biofilms by DCs, indicating IL-1β as pivotal cytokine induced by curli amyloid. A previous study established that curli activate the inflammasome in macrophages^41^, and our findings extend this paradigm to DCs and, crucially, identify curli as a direct ligand of caspase-11. Interestingly, curli rewrites the canonical two-signal requirement for inflammasome activation, because curli provides both priming and activation as a single stimulus, as evidenced by dose-dependent IL-1β secretion, caspase-1 cleavage, GSDMD cleavage, and ASC speck formation, none of which were further enhanced by prior LPS priming. This property implies that DCs encountering biofilm-derived curli at an infection site do not require prior inflammatory conditioning to mount a full inflammasome response.

To date, only two ligands, LPS ^31,32^ and OxPAPC^37^, have been shown by SPR to directly engage caspase-11; curli is now the third, and the first amyloid to do so. Its binding affinity (KD ≈ 5.69 × 10⁻⁷ M) is intermediate between LPS and OxPAPC, placing it in a biologically meaningful range: sufficient to activate caspase-11 and initiate non-canonical inflammasome signaling ^30,31^, yet calibrated to drive a non-lytic rather than fully cytolytic response. This discovery raises the broader question of whether caspase-11 and its human orthologs function as PRR for intracellular amyloids, a category encompassing other bacterial amyloids as well as endogenous aggregates such as amyloid-β, tau, α-synuclein, and SAA, all of which share the cross-β sheet architecture and have been linked to IL-1β production^71^.

We found that curli primes pro-IL-1β expression through a TLR2/TLR4–MyD88/TRIF signaling axis, with each adaptor making non-redundant contributions: MyD88-dependent signaling drives pro-IL-1β transcription, while TRIF-dependent IFN-β signaling is more selectively required for GSDMD cleavage, possibly through caspase-11 upregulation ^33^. Curli was previously shown to activate TLR2^15,41^, and its activation of TLR4 is a novel finding of this study.

Curli/TLR4/TRIF axis thus actively primes cytosolic non-canonical inflammasome activation by upregulation of caspase-11, highlighting an integrated innate response in which a single amyloid stimulus engages both surface and cytosolic signaling pathways. We confirmed that this TLR4 engagement was LPS-independent using polymyxin B neutralization and *Salmonella msbB* mutant, which produces hypo-stimulatory penta-acylated LPS ^52–56^ . Our results expand the known repertoire of TLR4 ligands, consistent with the reports that mammalian amyloids, including amyloid-β, also activate TLR2 and TLR4^72,73^. The same *msbB* mutant LPS has also been shown to not activate caspase-11^54^, excluding contaminant LPS as a potential confounding factor in the effects we attribute to curli.

To further establish curli specificity, we expressed and purified recombinant CsgA, fibrillized *in vitro*, and demonstrated that this form of *synthetic* curli alone fully recapitulate caspase-11 binding, inflammasome activation, and IL-1β release, validating curli as a bona fide non-canonical inflammasome ligand. The conservation of this response in hu-moDCs, including dose-dependent activation of caspase-5 (the human ortholog of murine caspase-11), confirms the pathway is not species-specific and strengthens translational relevance. As a biological caveat, LPS and curli co-exist within biofilms, and their cooperative or competitive recognition during infection by WT LPS-expressing biofilms remains possible and is the subject of ongoing investigation.

Mechanistically, curli fibers are phagocytosed and trafficked through the endolysosomal pathway, evidenced by curli colocalization with Rab5 and LAMP2. Upon cytoplasmic access, likely via the endolysosomal rupture evidenced by galectin-3 staining, curli engages caspase-11. This defines a hierarchical cascade: TLR2/TLR4 priming → endocytosis → endolysosomal escape → caspase-11 activation → GSDMD cleavage and pore formation→ K⁺ efflux → NLRP3– caspase-1 activation → mature IL-1β secretion ^30,32,34^. The selective suppression of IL-1β but not IL-6 by extracellular K⁺ supplementation confirms that TLR-driven priming and NLRP3 activation are separable. Notably, caspase-8 was not involved, positioning curli as a non-apoptotic, non-LPS activator of the non-canonical inflammasome.

Despite robust IL-1β secretion, curli-stimulated DCs do not undergo pyroptosis, exhibiting no LDH release, preserved ATP and mitochondrial potential, and sustained SYTOX Green exclusion over 22 hours. We propose that this reflects mechanistic constraints: curli must first escape endolysosomes via phagolysosomal rupture before engaging caspase-11, thus only a fraction of phagocytosed fibers reach the cytosol at any given time and engages caspase-11 with lower potency than transfected intracytoplasmic LPS ^30,31^. Moreover, mature GSDMD p30 is restricted to the plasma membrane, with no mitochondrial accumulation; its failure to engage the mitochondrial, cardiolipin driven amplification loop that escalates pore density under lytic conditions ^44^, is consistent with native PAGE showing only weak, late-forming GSDMD oligomers. Finally, NINJ1 fails to form the high-molecular-weight oligomers (>480 kDa) required for membrane rupture ^43^, providing a further downstream brake on lytic pore formation. Future studies will discover whether ESCRT-III membrane repair removes nascent pores before lytic thresholds are reached ^38,74^. A puzzle remains, about why pores permissive for IL-1β (17 kDa) secretion exclude SYTOX Green (∼600 Da) intake, which may reflect electrostatic filtering, as the GSDMD pore conduit is predominantly negatively charged and selectively favors passage of mature positively charged IL-1β while repelling negatively charged molecules of comparable size^75^. Rather than representing incomplete pyroptosis, DC hyperactivation is an actively safeguarded survival shaped by stimulus identity, constrained caspase-11 engagement, absent mitochondrial amplification, and possible membrane repair.

The functional consequence of preserved DC viability is immunologically significant. Curli-hyperactivated DCs show an approximately 5-fold increase in draining lymph node recovery compared to untreated cells, and a 2.4-fold increase over LPS-activated DCs, whereas pyroptotic DCs fail to migrate. By maintaining membrane integrity, mitochondrial potential and motility while upregulating co-stimulatory molecules and pro-inflammatory cytokines^6^, hyperactivated DCs may travel more as antigen-loaded APCs to sites of T cell priming sites, where the magnitude of T cell activation is directly proportional to the number of arriving DCs^76^, and improve host defense. An alternative interpretation warrants consideration, however: DCs that have internalized dispersed biofilm material could also disseminate biofilm components or viable bacteria to lymph nodes, a possibility that physiologically relevant infection models should address.

Collectively, these findings establish curli as a novel caspase-11 ligand and unique bacterial inducer of DC hyperactivation, engaging both canonical and non-canonical inflammasome pathways to drive potent cytokine secretion while preserving cell viability. Since curli is essential for biofilm structural integrity^70^, it plays a dual biological role: promoting bacterial persistence while simultaneously functioning as an immunostimulatory PAMP that licenses robust DC activation, precisely the kind of evolutionarily conserved molecular pattern that Charles Janeway envisioned as the initiator of innate immunity.

## Limitations of this study

Some limitations merit acknowledgment. We treated DCs with curli and biofilm *in vitro,* and whether DCs recognize intact mature biofilms *in vivo* or respond to curli on immature biofilms prior to full maturation, remains to be determined. Moreover, future work will define how curli-driven DC hyperactivation shapes adaptive immunity during chronic infection and explore whether other bacterial or endogenous amyloids engage the same caspase-11-NLRP3 axis. These investigations may reveal therapeutic targets for disrupting biofilm persistence, enhancing immune clearance, and improving outcomes in patients with biofilm-associated infections.

## Acknowledgements

We thank Dr. Shruti Choudhary (Dr. Paul Thompson’s lab), at UMass Chan SPR Core facility for her assistance with the caspase 11/curli SPR binding experiments. We thank Drs. Douglas T. Golenbock and Kenneth Rock for providing transgenic mice.

## Authors contributions

SM, KAF and SG designed the project. SM performed most of the experiments. GH designed and performed the CsgA purification. LMW generated E. coli biofilms. GH and BL performed animal studies. KAF supervised and SM and AM performed confocal microscopy studies. EL supervised and AD performed Ninj1 studies. VAR provided expertise in caspase-11 and supervised PS who bred Caspase 11 mice and harvested bones for *in vitro* studies. CT supervised and KG generated Salmonella biofilms and purified curli. SM and SG wrote the manuscript. KAF and SG co-mentored SM. SG supervised all research. All authors read and approved the manuscript.

This work was supported by NIH R03AI173522-01 and R21AI180971-01 (to SG), NIH R01 AI153325, R01AI171568, and R21AI180971-01 (ÇT), NIH R21 AI182982 and RO1AI181309 and The Research Council of Norway, Center of Excellence funding scheme project 223255/F50 (EL).

## Declaration of generative AI and AI-assisted technologies in the manuscript preparation process

During the preparation of this work, the author(s) used Claude (Anthropic) to assist with language editing and style. The author(s) reviewed and edited the output as needed, and take full responsibility for the content of the published article.

## Materials and Methods

### Ethics

*Mice.* Experimental mice and breeding mice were housed in the same BSL1 room maintained in the UMass Chan Medical School animal facility, an American Association for the Accreditation of Laboratory Animal Care–accredited facilities. Studies were approved by UMass Chan Institutional Animal Care and Use Committee (IACUC).

*Human cells.* De-identified human blood leukopaks were obtained from UMass Memorial Hospital under approval from the Institutional Review Board of the University of Massachusetts Chan Medical School. All donors provided written informed consent. All studies were approved and monitored by the Institutional Biosafety Committee at the University of Massachusetts Chan Medical School.

### Mice strains

WT C57BL/6 mice, *Tlr2* KO^77^, *Tlr4* KO ^78^ were purchased from Jackson Laboratory. We crossed *Tlr2* KO and *Tlr4* KO to make *Tlr2/4* double KO, *Asc* KO ^79^, *Nlrp3* KO ^80^, *Caspase 1* KO ^81^, *Trif* KO^82^, *Myd88* KO ^83^, *Caspase 1/8/11/Ripk3* quadruple KO ^84^, *Caspase 8/Ripk3* double KO ^62^, *Caspase 1/11* double KO ^85^ and *Gsdmd* KO ^34^ mice were bred at UMass Chan medical School. 8-16 weeks old, were used for all experiments. *Caspase 11* KO mice ^86^ were bred at UConn health and bones from these mice were sent for isolation and differentiation. All mice were 8–16 weeks of age at the time of experiments.

### Reagents

The following reagents were used: FBS (GeminiBio, Cat# 100-106-500); IMDM (Sigma, Cat# 51471C); phenol red–free IMDM (Gibco, Cat# 21056023); recombinant mouse GM-CSF (BioLegend, Cat# 576308); penicillin-streptomycin (Gibco, Cat# 10378016); Lipofectamine 2000 (Invitrogen, Cat# 11668019); LPS *E. coli* O111:B4 (Sigma, Cat# L2630); LPS-EB (InvivoGen, Cat# tlrl-pelps); LPS-EB Biotin (InvivoGen, Cat# tlrl-lpsbiot); Pam3CSK4 (InvivoGen, Cat# tlrl-pms); Pam3CSK Biotin (InvivoGen; Cat# tlrl-bpms); nigericin (Sigma, Cat# N7143); R848 (InvivoGen, Cat# tlrl-r848-1); MCC950 (InvivoGen, Cat# inh-mcc); z-VAD-FMK (Santa Cruz, Cat# sc3067); polymyxin B (Sigma, Cat# tlrl-pmb); TAK-242 TLR4 inhibitor (Sigma, Cat# 614316); and KCl solution (Sigma, Cat# 60142).

### Generation of murine bone marrow-derived Dendritic Cells (muDCs)

Bone marrow-derived dendritic cells were generated from 8–12-week-old mice as previously described ^45,87^. Femurs and tibiae were dissected under sterile conditions, cleaned, sterilized briefly in 70% ethanol, air dried and bone marrow was flushed three times per bone using a 27G needle. Cell suspensions were dissociated, filtered through sterile nylon mesh, and centrifuged at 1,500 rpm for 10 min. Cells were resuspended in DC differentiation medium (complete IMDM supplemented with recombinant GM-CSF) at 1 × 10^6^ cells/mL and plated at 1 mL/well. On day 3, equal volumes of fresh medium were added per well. On days 5 and 6, half the medium was replaced without disturbing semi-adherent cells. muDCs were used between days 7 and 9. Typical yield was ∼0.75–1 × 10^6^ cells/well with >80% CD11c positivity by flow cytometry.

### Generation of Human Monocyte-Derived Dendritic Cells (hu-moDCs)

PBMCs were isolated from de-identified healthy donor leukopaks by Ficoll-Paque PLUS density centrifugation (400 × g, 30 min, room temperature, brake off; GE Healthcare; Cat# 7-1440-02). Monocytes were enriched by positive immunomagnetic selection using the EasySep Human CD14 Positive Selection Kit II (StemCell Technologies, Cat# 17858). Purified monocytes were plated at 1 × 10^6^ cells/mL in complete RPMI 1640 supplemented with 10% heat-inactivated FBS, 1% penicillin-streptomycin, 2 mM L-glutamine, 50 ng/mL human GM-CSF (BioLegend, Cat# 572904), and 50 ng/mL human IL-4 (BioLegend, Cat# 574006) as previously described with some modifications^48^. On day 3, half the medium was replaced with fresh cytokine-supplemented medium without disturbing the semi-adherent cells. On day 7, half of the medium was changed and then DCs were stimulated with indicated ligands for indicated time points.

### Curli purification from Salmonella Biofilms

Curli was purified from the *S*. Typhimurium IR715 *msbB* mutant as previously described with modifications^6,88^. Bacteria grown in liquid YESCA medium supplemented with 4% DMSO at 26°C for 72 h with shaking (200 rpm). Curli was purified from bacterial pellets by sequential nuclease, lysozyme, and SDS treatments followed by sonication, as previously described ^89^, with additional nuclease digestion and washing steps to remove contaminating cellular components. Purified curli aggregates were isolated by SDS-PAGE, extracted with ethanol, resuspended in sterile water, and quantified by BCA assay. Final preparations were adjusted to a protein concentration of 1 mg/ml.

### Recombinant CsgA Expression and Purification

Recombinant *Salmonella* CsgA was expressed and purified as adapted from Perov et al. (2019)^90^. Briefly, plasmid encoding CsgA with a C-terminal His_6_ tag (cloned into pET21d; GenScript) was transformed into *E. coli* BL21 cells. Overnight cultures in LB supplemented with 50 µg/mL carbenicillin and 1% glucose were diluted 1:100 into the same medium and grown at 37°C with shaking at 140 rpm to OD_600_ 0.8–0.9, then induced with 0.5 mM IPTG for 1 hr. Cells were harvested at 6,000 × g for 15 min and stored at −80°C. Pellets were resuspended in 25 mL lysis buffer (8 M guanidinium hydrochloride), sonicated, and clarified by centrifugation. The lysate was incubated with His-Pur Ni-NTA resin (Thermo Fisher, Cat# 88221) pre-equilibrated in lysis buffer for 2 hr at room temperature. After washing with ice-cold PBS and PBS containing 25 mM imidazole, protein was eluted with 250 mM imidazole and desalted into PBS. Thus purified CsgA was subjected to a single freeze–thaw cycle, a procedural step that we discovered promotes amyloid fibril formation. Fibrillated CsgA was pelleted at 15,000 × g for 15 min at 4°C and washed three times with cell culture-grade DPBS to reduce residual endotoxin contamination.

### Thioflavin (ThT) Staining

Amyloid content of curli and CsgA fibers preparations was confirmed using Thioflavin T (ThT) fluorescence, as previously described ^64^. Curli and CsgA fibers were incubated with ThT (5 µM final concentration) in black 96-well plates or biofilms on agar plates at room temperature for 10 min. Fluorescence was measured on a SpectraMax plate reader (excitation 440 nm, emission 500 nm). Fibril particle size and morphology were assessed by brightfield and ThT fluorescence imaging on an Agilent BioTek Lionheart LX automated microscope.

### Generation of UTI-89 biofilms

Uropathogenic *E. coli* (UPEC) UTI89 biofilms were generated as follows. Overnight planktonic cultures in LB broth (37°C, 200 rpm) were diluted 1:100 into YESCA medium, plated at 2 mL/well in 24-well plates, and incubated for 72 hr at 28°C under static conditions on a slight incline. Planktonic bacteria were removed by washing the biofilms three times with PBS, then the biofilms were resuspended in 200–250 µL/well PBS supplemented with gentamicin (50 µg/ml) to inhibit bacterial growth, pooled, disaggregated mechanically by vigorously shacking the tubes, and O.D. was adjust to 0.5 at 600nm. Biofilm mass was quantified by BCA and O.D. at 600nm and expressed as Equivalent Colony Forming Unit (eCFU). To decrease the contribution of LPS to biofilm-mediated DC activation, UTI89 *msbB* mutant, producing a pentacylated Lipid A that is unable to activate human TLR4 and is a weak agonist of murine TLR4^52,53^ (generated by Dr. Amir Mitchell’s lab, UMass Chan Medical School, manuscript in preparation), was used. To image the morphology and curli expression, biofilms were generated on YESCA plates ^91^ and left unstained or stained with Congo Red (500 µg/mL) and Coomassie Bright Blue (100 µg/mL) for 20 minutes, and then gently wash. Congo Red is a dye that intercalates into amyloid cross-β-sheet structures and serves as a colorimetric indicator of curli production in Enterobacteriaceae ^92^, Coomassie Bright Blue stains proteins in general. Unstained and CR/CB-stained biofilms were photographed using a smartphone camera with Macro Control enabled (iPhone 14 Pro Max, Apple Inc., Cupertino, CA, USA). Amyloid expression was also detected by staining colony biofilms with ThT, as described above, and biofilms were imaged using a LiCor ODYSSEY M system.

### Immunoblot analysis

Cells were stimulated in IMDM supplemented with 5% FBS for the specified time points. For combined supernatant and lysate preparations, 50 µL of lysis buffer [10% Triton X-100, 10 mM DTT, protease inhibitor cocktail (Sigma, Cat# 11836153001)] was added to samples, followed by 120 µL 4× loading dye (8% SDS, 400 mM DTT, 0.4% bromophenol blue, 40% glycerol, 200 mM Tris pH 6.8). Cell lysates alone were prepared by direct lysis in 2× LDS sample buffer (Thermo Fisher, Cat# AAJ61942AD). All samples were heated at 95°C for 15-20 min.

Proteins were resolved on 10–13% SDS-PAGE gels and transferred onto 0.45 µm nitrocellulose membranes (Bio-Rad, Cat# 1620115) using the Trans-Blot Turbo system (2.5 A, 25 V, 10 min). Membranes were blocked in 5% skimmed milk in TBST for 1 hr at room temperature, then incubated overnight at 4°C with primary antibodies in 5% BSA/TBST. After washing in TBST, membranes were incubated with HRP-conjugated secondary antibodies (1:5,000 in 5% BSA/TBST) for 2 hr at room temperature. Signal was detected with Clarity Western ECL Substrate (Bio-Rad, Cat# 1705061) on a ChemiDoc Imaging System.

Primary antibodies: anti-caspase 1 (AdipoGen, Cat# AG-20B-0042); anti-IL-1β (CST, Cat# 8689); anti-NLRP3 (AdipoGen, Cat# AG-20B-0014); anti-ASC (AdipoGen, Cat# AG-25B-0006); anti-GSDMD (Abcam, Cat# ab209845); anti-caspase 11 (Abcam, Cat# EPR18628); anti-NINJ1 (clone 25, kind gift of Genentech); anti-β-actin (Sigma, Cat# A2228); anti-IκBα (CST, Cat# 9242); anti-phospho-IκBα (CST, Cat# 2859); anti-NF-κB p65 (CST, Cat# 8242S); anti-phospho-NF-κB p65 Ser536 (CST, Cat# 3033); anti-caspase 8 (Enzo, Cat# ALX-804-447); anti-caspase 5 (CST, Cat# 4429); anti-caspase 3 (CST, Cat# 9662).

### Blue Native-Page Gel

After stimulation, the supernatant was removed from muDCs, and cells were gently washed once in 1x PBS before lysis at indicated time points. Cells were harvested with Native lysis buffer consisting of 1% Digitonin (Sigma D141), 50mM Tris pH7.4, 150mM NaCl, and 1x Protease inhibitor cocktail (Roche). Lysates were then centrifuged at 20,000g, 4C, 30min, and mixed with NativePAGE sample buffer 4x (ThermoFisher Scientific). Lysate samples were quantified by Bradford assay (Bio-Rad) and equated for gel loading. For analyzing NINJ1 polymerization, the Genetech NINJ1 antibody (clone 25) ^43^ was used, and bands above 480 kDa were considered positive for NINJ1 polymerization and activation.

### Enzyme-Linked Immunosorbent Assay (ELISA)

Cytokine concentrations in cell-free supernatants were quantified by sandwich ELISA using DuoSet kits (R&D Systems) for murine IL-1β (Cat# DY401), murine IL-6 (Cat# DY406), human IL-1β (Cat# DY201), and human IL-6 (Cat# DY206) according to manufacturer’s instructions. Absorbance was read at 450 nm (reference 570 nm) on a SpectraMax plate reader. Concentrations were interpolated from the standard curve and expressed as pg/mL.

### Immunofluorescence staining

muDCs were seeded at 3 × 10^5^ cells/well in 4-chamber slides (Thermo Fisher, Cat# 154917PK). Following stimulation, cells were washed three times with PBS, fixed with 4% PFA for 15 min at room temperature, and blocked/permeabilized with 10% normal goat serum plus 0.1% saponin for 1 hr. Cells were incubated overnight at 4°C with primary antibodies: rabbit anti-ASC (1:200 dilution, AdipoGen, Cat# AG-25B-0006-C100), mouse anti-Rab5 (1:500 dilution, Santa Cruz, Cat# sc-46692), rat anti-LAMP2 (1:500 dilution, Abcam, Cat# ab13524), or mouse anti-Galectin-3 (1:500 dilution Abcam, Cat# ab2785). Rab5 and LAMP2 served as markers of early endosomes and lysosomes, respectively, to assess endolysosomal trafficking of curli following phagocytic uptake. For co-localization studies, curli was pre-labeled with Congo Red (10 µg/mL). Galectin-3 was used as a marker of lysosomal membrane rupture. Secondary antibodies: species-matched Alexa Fluor 488-conjugated antibodies (Thermo Fisher, Cat# A-11008, A-11006, A-11001) were applied for 1 hr at room temperature. Slides were mounted in DAPI-containing medium (Vector Laboratories, Cat# H-1200). Imaging was performed on a Leica SP5 confocal microscope. For ASC speck quantification, a minimum of 150 cells per condition were counted across at least three independent fields of view.

### Lactate Dehydrogenase (LDH) Cytotoxicity Assay

Plasma membrane rupture was assessed by measuring LDH release using the CytoTox 96 Non-Radioactive Cytotoxicity Assay (Promega, Cat# G1780). Briefly, both murine and human DCs were stimulated in phenol red–free IMDM with 5% FBS. Cell-free supernatants (50 µL) were transferred to 96-well plates, mixed with an equal volume of substrate, and incubated at room temperature for 10 min protected from light. The reaction was stopped upon color change to dark pink. Maximum LDH release was determined from cells lysed with the kit lysis solution. Absorbance was measured at 490 nm on a SpectraMax plate reader. Percent cell death = (sample absorbance / maximum release absorbance) × 100.

### CellTiter-Glo Luminescent Cell Viability Assay

Intracellular ATP levels were measured as an orthogonal viability readout using the CellTiter-Glo Luminescent Cell Viability Assay (Promega, Cat# G7570). An equal volume (200 µL) of reconstituted CellTiter-Glo reagent was added to each well containing 200 µL of culture medium. Contents were mixed for 2 min on an orbital shaker and incubated for 10 min at room temperature to stabilize the luminescent signal. Luminescence was recorded in relative light units (RLU) on a SpectraMax plate reader. Reduced RLU relative to untreated controls indicates cell death.

### Live-Cell Imaging of Cell Death Kinetics

Cell death kinetics were monitored by live-cell imaging using the membrane-impermeant dye SYTOX Green (Thermo Fisher, Cat# S7020). muDCs were stimulated with curli (10 µg/mL) or left untreated for 16 hr prior to adding dyes and start imaging. This extended stimulation period before adding the live dye accommodates the phagocytic nature of DCs and allows curli uptake and inflammasome activation without confounding non-specific dye uptake during stimulation. Pyroptosis controls were primed with LPS (100 ng/mL, 2–3 hr) and then nigericin (10 µM) was added immediately before imaging. Cells were stained with Hoechst 33342 (InvivoGen, Cat# 34580; 4 µM) and SYTOX Green (0.5 µM) for 10 min at 37°C prior to imaging. Live-cell imaging was performed on an Agilent BioTek Lionheart FX automated microscope (37°C, 5% CO2), acquiring fluorescence images every 20 min over 6–8 hr. Cellular objects <40 µm (Hoechst channel) were excluded from analysis. Cell death was expressed as the ratio of SYTOX Green to Hoechst mean fluorescence intensity, normalized to untreated controls at time zero, multiplied by 100.

### Subcellular Protein Fractionation

Cytoplasmic and membrane fractions were isolated using the Subcellular Protein Fractionation Kit for Cultured Cells (Thermo Fisher, Cat# 78840), and mitochondrial fractions using the Mitochondria Isolation Kit for Cultured Cells (Thermo Fisher, Cat# 89874), per manufacturer instructions. For each condition, 10 × 10^6^ muDCs were seeded in 6-well plates, stimulated, and detached by scraping. Cell pellets were washed with ice-cold PBS prior to fractionation.

For cytoplasmic and membrane fractions, cells were resuspended in Cytoplasmic Extraction Buffer (CEB) supplemented with protease inhibitors, incubated on ice for 10 min, and centrifuged at 500 × g (4°C, 5 min) to pellet nuclei. The supernatant was re-centrifuged at 3,000 × g to yield the cytoplasmic fraction. The remaining pellet was resuspended in Membrane Extraction Buffer (MEB) supplemented with protease inhibitors and centrifuged at 3,000 × g (4°C, 5 min) to yield the membrane fraction. For mitochondrial isolation, cells were resuspended in Reagent A (with protease inhibitors), then Reagent B was added and the suspension vortexed briefly to disrupt the plasma membrane. After centrifugation at 700 × g (4°C, 10 min) to remove nuclei and debris, mitochondria were pelleted at 3,000 × g for 15 min, washed with Reagent C, and resuspended in RIPA buffer with protease inhibitors. Fractionation quality was confirmed by immunoblotting for calreticulin (ER/membrane; Thermo Fisher, Cat# MA5-32131), caspase-1 (cytoplasm; AdipoGen, Cat# AG-20B-0042), and COX IV (mitochondria; Thermo Fisher, Cat# MA5-15078). GSDMD distribution across fractions was assessed using anti-GSDMD (Abcam, Cat# EPR19828).

### Co-Immunoprecipitation (Co-IP) Assay

Co-IP was adapted from previously described protocols ^32,37^. muDCs (1 × 10^7^/condition) were primed with LPS (100 ng/mL, 3 hr) or curli (10 µg/mL, 3 hr) to induce caspase-11 expression. Curli-primed cells were further stimulated with or without biotinylated curli (50 µg/mL) for an additional 6 hr. Cells were washed with ice-cold PBS, scraped, pelleted (500 × g, 5 min, 4°C), and lysed for 15 min in lysis buffer [50 mM Tris-HCl pH 7.6, 150 mM NaCl, 2 mM EDTA, 1% Triton X-100, protease inhibitor cocktail (Roche, Cat# 11836153001)] followed by centrifugation at 13,000 rpm for 30 min at 4°C. Input controls (50 µL) were reserved and stored at −80°C. Pre-cleared lysates were incubated overnight at 4°C with rotation with 40 µL Protein A/G magnetic beads plus biotinylated curli, LPS, or Pam3CSK4 (samples stimulated with both curli and biotinylated curli received beads alone). Curli was biotinylated using EZ-Link™ NHS-PEG4-Biotin, No-Weigh™ Format (Thermo A39259). Biotinylated LPS EB-biotin was purchased from Invivogen (tlrl-lpsbiot). Beads were washed three times with lysis buffer, and immune complexes were eluted in 100 µL 2× LDS buffer (95°C, 10 min). Eluates were analyzed by SDS-PAGE and immunoblotted for caspase-11 and caspase-3 to confirm co-precipitation.

### Surface plasmon resonance (SPR)

SPR experiments were performed to assess direct binding between caspase-11 and curli, using methods adapted from Shi et al. ^32^ Briefly, recombinant murine caspase-11 was immobilized on a sensor chip, and LPS, curli, CsgA, MDP, or Pam3CSK4 were flowed over the chip as analytes at a range of concentrations to determine binding kinetics and equilibrium dissociation constants (K_D_). LPS served as a positive control, given its established direct, high-affinity interaction with caspase-11 via its CARD domain^32^, while MDP and Pam3CSK4, ligands for NOD2 and TLR2 respectively that are not known to engage caspase-11 directly, served as negative controls. Curli and CsgA showed dose-dependent, saturable binding to immobilized caspase-11, whereas MDP and Pam3CSK4 produced no specific binding signal above background.

### *In Vivo* Dendritic Cell Migration Assay

muDCs were labeled with Cell Trace Violet (CTV; Thermo Fisher, Cat# C34557) to track donor cells in recipient lymph nodes. muDCs were resuspended in CTV/PBS, incubated at 37°C for 20 min in the dark, quenched with 5 volumes of complete IMDM, and returned to culture overnight to allow muDC to rest. Labeled muDCs were then stimulated with curli (10 µg/mL) or LPS (100 ng/mL) for 15 hr. Pyroptosis controls were primed with LPS (100 ng/mL, 2 hr) followed by nigericin (10 µM) prior to injection. Untreated CTV-labeled muDCs served as negative migration controls.

After stimulation, cells were washed twice with sterile PBS and viability assessed by trypan blue exclusion. For each condition, 2 × 10^6^ CTV-labeled muDCs in 40 µL sterile PBS were injected subcutaneously into the hind footpad of syngeneic C57BL/6 recipient mice (n = ∼5/group). After ∼36 hr, mice were euthanized, popliteal lymph nodes harvested, and donor DC recovery analyzed by flow cytometry on a Cytek Aurora. The absolute number of injected CTV^+^ cells recovered in draining lymph nodes served as the primary measure of migratory capacity. Data represent the combined analysis of five independent experiments.

### Statistical Analysis

Statistical analyses were performed using GraphPad Prism v10.0. Data are presented as mean ± SEM of at least three independent biological replicates. Comparisons across three or more groups were performed by one-way ANOVA with Tukey’s, Dunnett’s, or Šídák’s post-hoc test as appropriate. Two-group comparisons used a two-tailed unpaired Student’s t-test. Two independent variables were assessed by two-way ANOVA with appropriate post-hoc correction. The specific test for each experiment is indicated in the corresponding figure legend. Significance thresholds: NS, not significant; *p < 0.05; **p < 0.01; ***p < 0.001; ****p < 0.0001.

## Supplemental Figures

**Figure S1.**
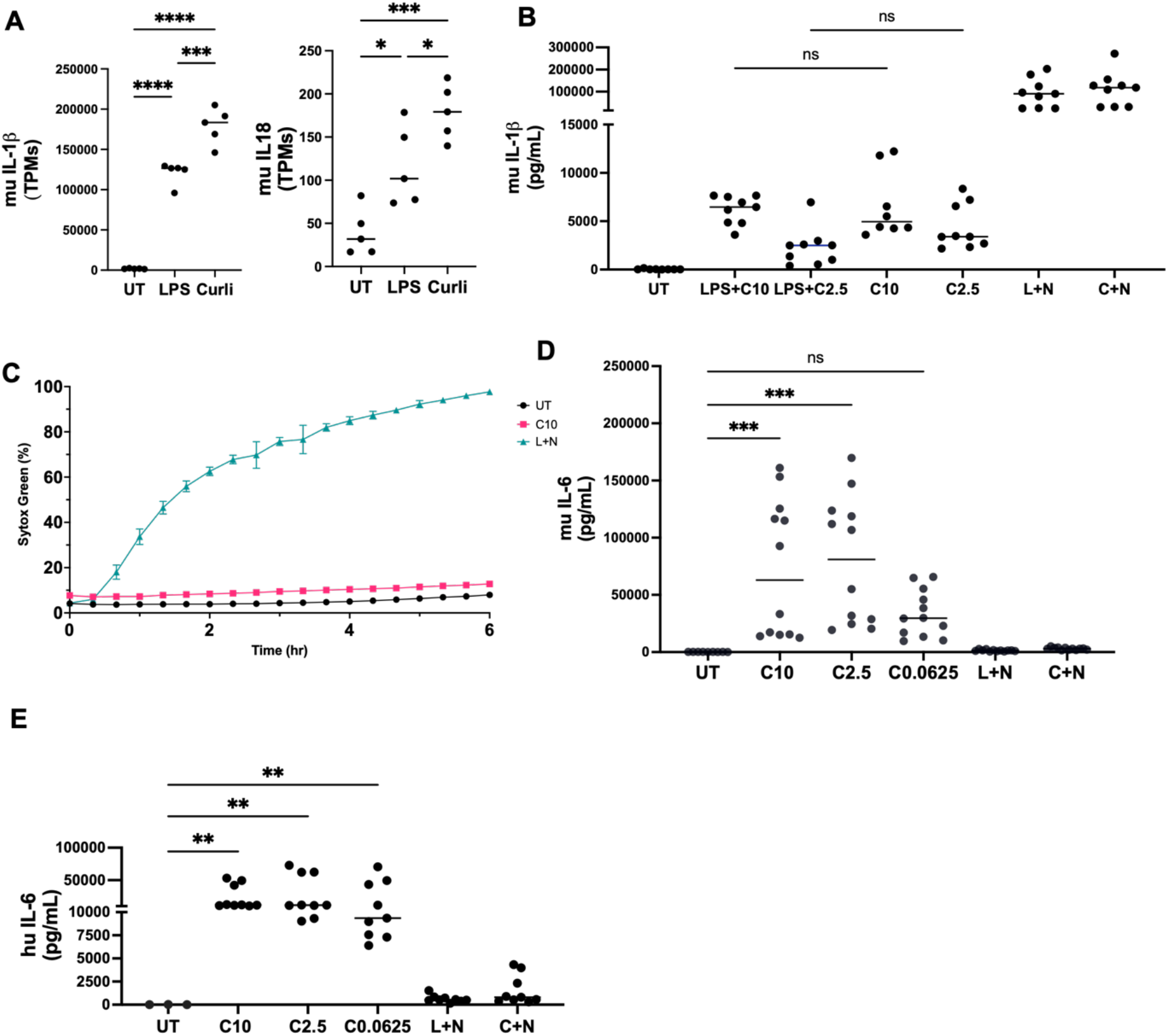
Curli activates the inflammasome without inducing cell death. (A*) Il1b* (*left*) and *Il18* (*right*) levels of transcript per million (TPMs) from RNA-seq analysis of murine WT DCs stimulated with LPS or curli for 6 hr compared to untreated controls. **(B)** Murine IL-1β release measured by ELISA in supernatants of WT DCs (20-22hrs) stimulated with 100 ng/ml of LPS combined with curli at 10 and 2.5 µg/mL (LPS+C10, LPS+C2.5), curli alone (10, 2.5 µg/mL), or with LPS and then Nigericin (10 µM)(L+N) or curli 10 µg/mL and then Nigericin (C+N) (3+3 hrs). **(C)** Real-time kinetics of plasma membrane permeabilization in murine WT DCs monitored by SYTOX Green uptake over 6 hr by live-cell imaging on a Lionheart automated microscope. Untreated (UT, black), curli 10 µg/mL (C10, pink), LPS + nigericin (L+N, teal). For curli-treated conditions, SYTOX Green was added 16 hr after the beginning of the stimulation to avoid non-specific dye uptake via phagocytosis; for L+N, SYTOX Green was added immediately following nigericin stimulation. Data expressed as percentage SYTOX Green-positive cells normalized to Hoechst-positive cells. **(D)** Murine IL-6 release measured by ELISA in supernatants of WT DCs stimulated for 22 hrs (curli) or 6 hours (L+N)(C+N) as indicated. **(E)** Human IL-6 release measured by ELISA in supernatants of hu-moDCs stimulated with curli at indicated concentrations (10, 2.5, 0.0625 µg/mL) for 22 hrs (curli) or 6 hours L+N, or C+N. Data represent mean ± SEM from at least three independent DC cultures (A), (B), and (D), and three independent experiments with three human donors (C). Statistical significance was determined by one-way ANOVA with Šídák’s or Dunnett’s multiple comparisons post-hoc test. *p < 0.05, **p < 0.01, ***p < 0.001, ****p < 0.0001; ns, not significant.

**Figure S2.**
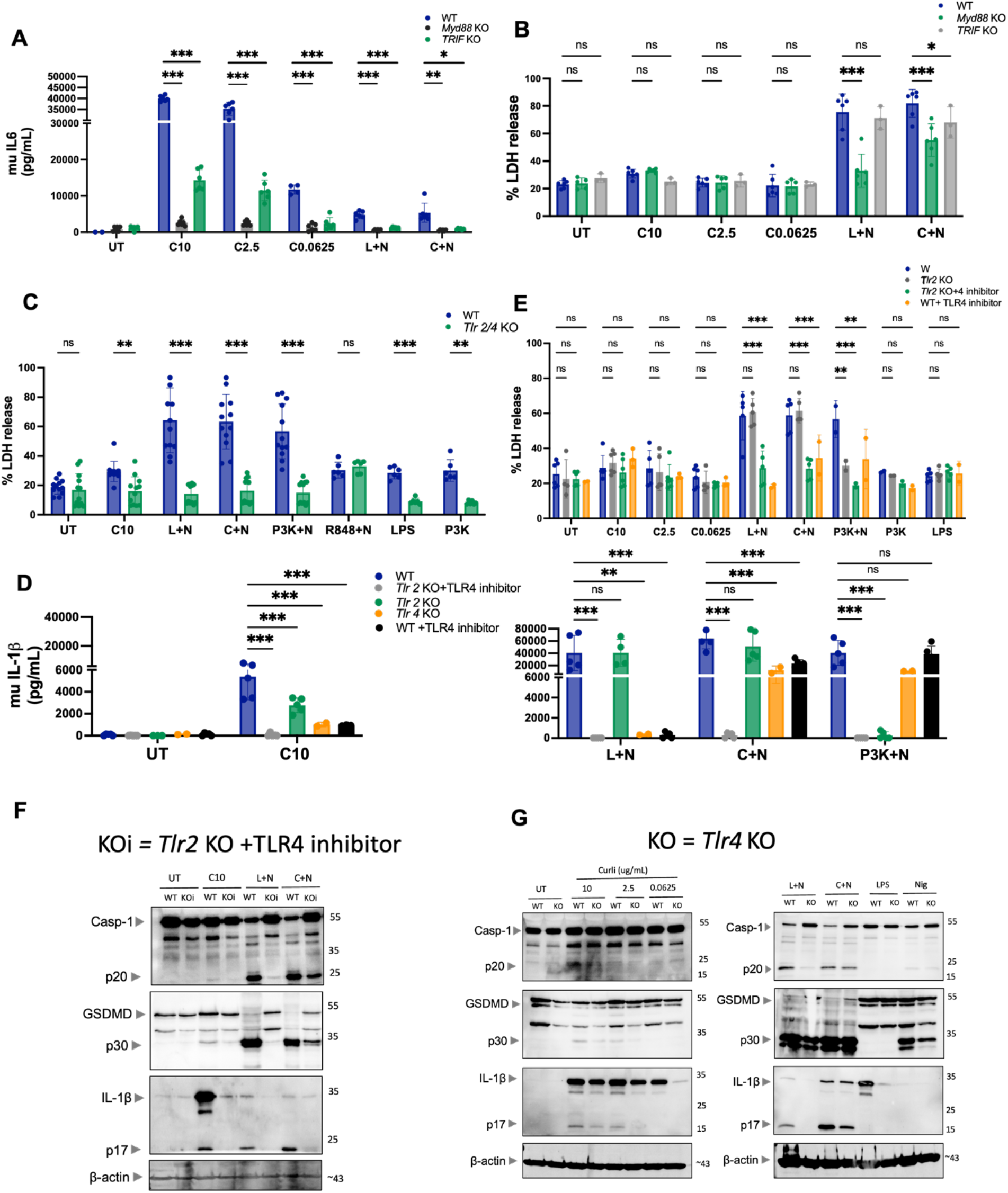
Curli primes the inflammasome through MYD88 and TRIF triggered by TLR2 and TLR4. **(A**) Murine IL-6 release measured by ELISA in WT (blue), *Myd88* KO (black), and *Trif* KO (green) muDCs stimulated with curli at the indicated concentrations for 20–22 hr, and L+N, or C+N for 5.5 hrs. **(B)** Cell death assessed by LDH release (% of maximum) in WT (blue), *Myd88* KO (black), and *Trif* KO (green) muDCs stimulated with curli at the indicated concentrations for 20–22 hr, and L+N, or C+N for 5.5 hrs. **(C)** Cell death assessed by LDH release (% of maximum) in WT (blue) and *Tlr2/4* DKO (green) muDCs stimulated with curli (C10), LPS, or P3K alone for 20–22 hr, and L+N, C+N, P3K+N, R848+N for 5.5 hrs. **(D)** Murine IL-1β release measured by ELISA in WT (blue), *Tlr2* KO + TLR4 inhibitor TAK-242 (grey), *Tlr2* KO (green), *Tlr4* KO (orange), and WT + TLR4 inhibitor TAK-242 (black) muDCs stimulated with curli (C10) alone (*left*) or with L+N, C+N, or P3K+N (*right*), demonstrating individual and combined contributions of TLR2 and TLR4 to inflammasome priming. TAK-242 (200 nM) was added 1 hr prior to ligand stimulation. **(E)** Cell death assessed by LDH release (% of maximum) in WT (blue), *Tlr2* KO (grey), *Tlr2* KO + TLR4 inhibitor TAK-242 (green), and WT + TLR4 inhibitor (orange) muDCs stimulated with the indicated conditions. **(F, G)** Immunoblot analysis of caspase-1 (pro-form and p20 cleavage fragment), GSDMD (pro-form and p30 cleavage fragment), IL-1β (pro-form and mature p17 fragment), and β-actin in WT and *Tlr2*KO + TLR4 inhibitor (KOi) muDCs **(F)** and in WT and *Tlr4* KO muDCs **(G)** stimulated with curli (C10) for 20–22 hr, and L+N, or C+N for 5.5 hrs. Data represent mean ± SEM from at least three independent biological replicates. Statistical significance was determined by multiple unpaired t-tests with Šídák’s correction for multiple comparisons between WT and KO or inhibitor-treated conditions within each treatment group. *p < 0.05, **p < 0.01, ***p < 0.001, ****p < 0.0001; ns, not significant.

**Figure S3.**
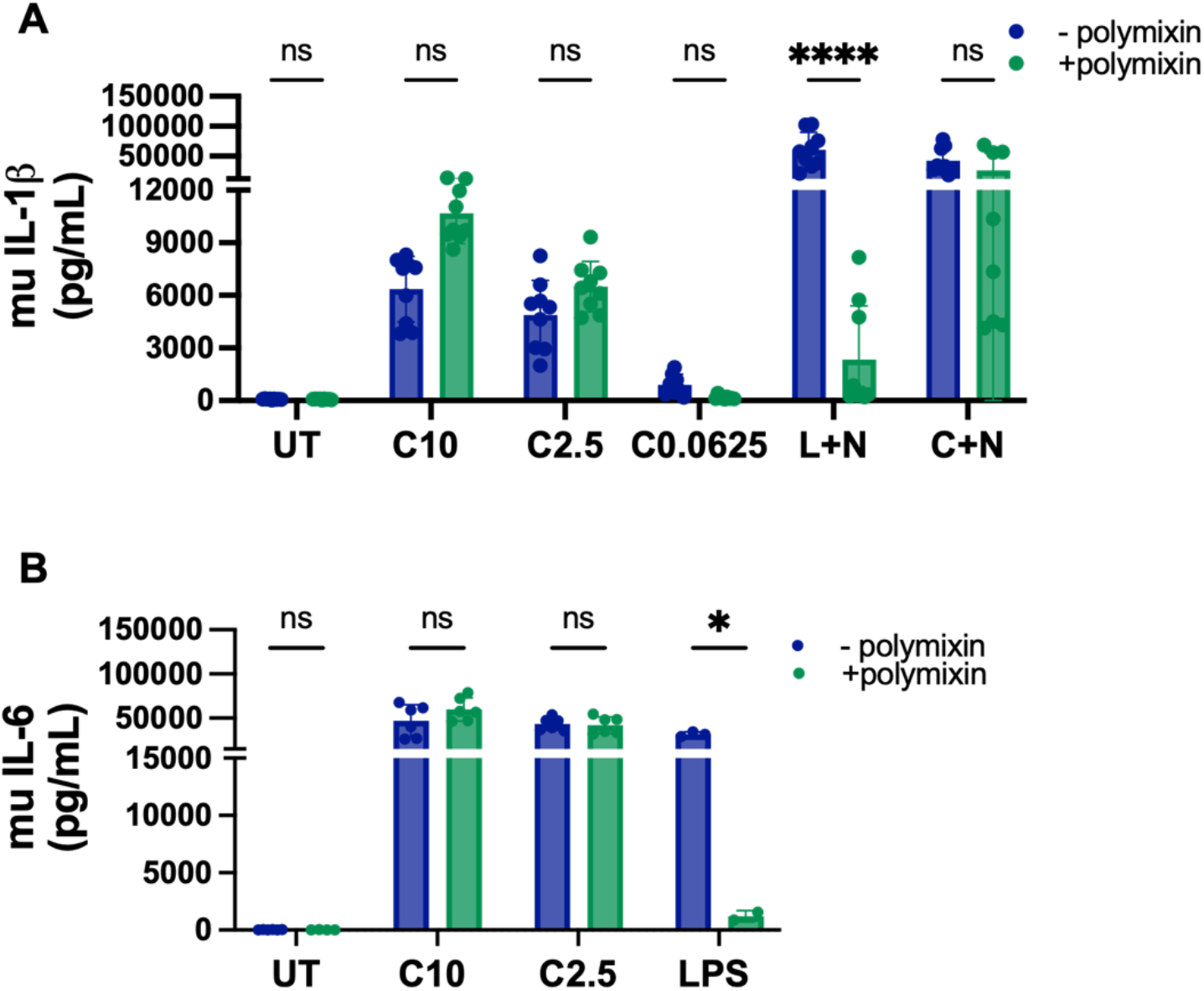
Curli-induced inflammasome activation is independent of LPS contamination. **(A)** IL-1β release and **(B)** IL-6 release, measured by ELISA in muDCs stimulated with curli at the indicated concentrations (C10, C2.5, C0.0625 µg/mL; 22 hours), L+N, or C+N (3+2.5 hours)in the presence (green) or absence (blue) of polymyxin B (10 µg/mL; added 1 hour before ligand). LPS served as a positive control for polymyxin B-mediated neutralization. UT, untreated; L+N, LPS + nigericin; C+N, curli (10 µg/mL) + nigericin. Data represent mean ± SEM. Statistical significance was determined by unpaired t-tests with Šídák’s correction for multiple comparisons *p<0.05, **p<0.01, ***p<0.001, ****p<0.0001; ns, not significant. Unless otherwise indicated, statistical comparisons were made between polymyxin B-treated and untreated conditions within each stimulation group. Data are representative of at least two independent experiments.

**Figure S4.**
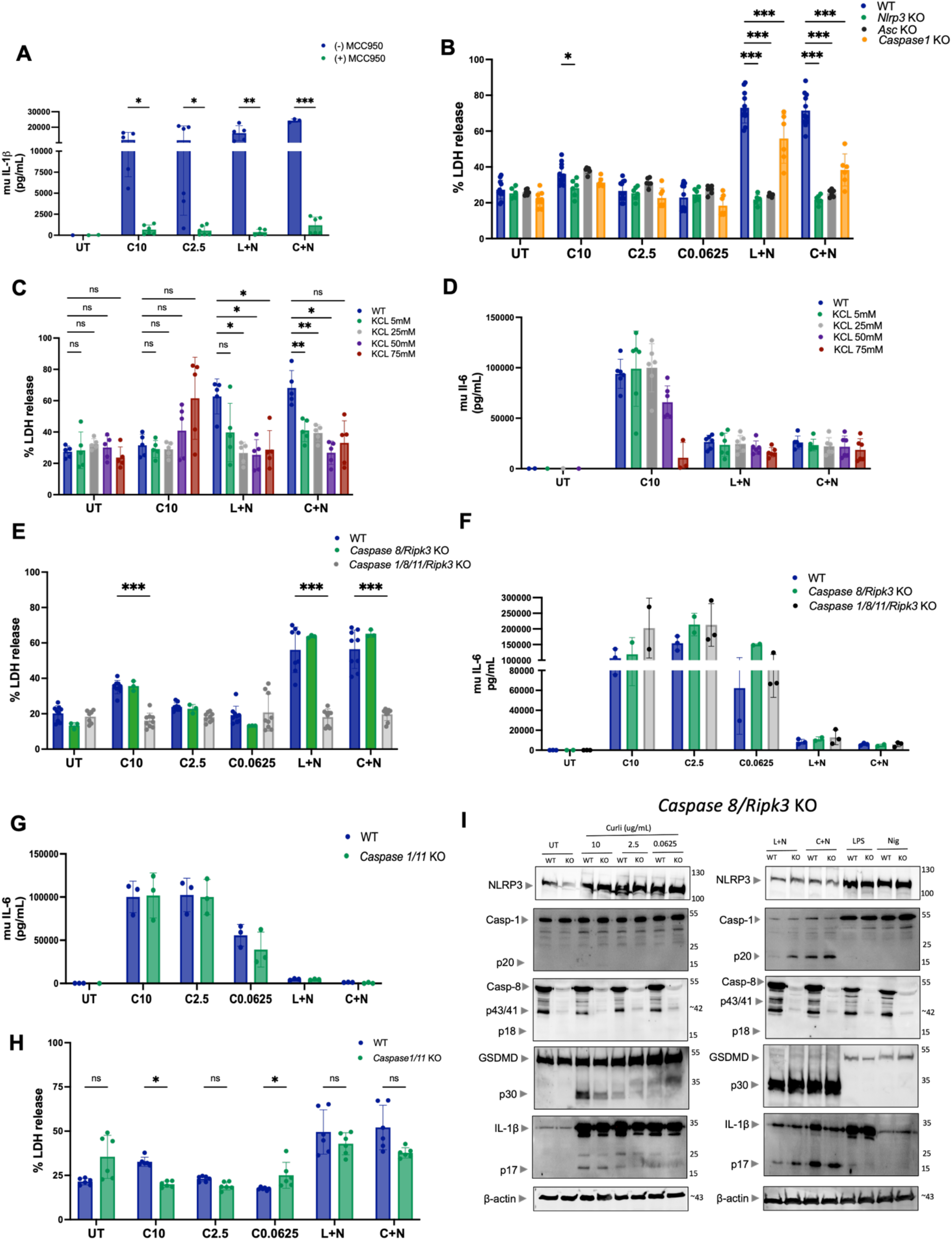
Curli-induced inflammasome activation is dependent on caspases 1/11 and NLRP3. **(A)** Murine IL-1β release by ELISA in WT muDCs stimulated with curli (C10, C2.5), L+N, or C+N ± MCC950 (10 µM, added 1 hr prior to stimulation). **(B)** Cell death (% LDH release) in WT, *Nlrp3* KO, *Asc* KO, and *Caspase1* KO muDCs under the indicated conditions. **(C)** Cell death (% LDH release) in WT muDCs pre-treated with extracellular KCl (5–75 mM) prior to stimulation with curli (C10), L+N, or C+N. **(D)** Murine IL-6 release measured by ELISA in KCl pre-treated muDCs under the indicated conditions, confirming that K⁺ efflux blockade does not impair TLR-dependent priming. **(E)** Cell death (% LDH release) in WT, *Caspase 8/Ripk3* DKO, and *Caspase 1/8/11/Ripk3* KO muDCs under the indicated conditions. **(F)** Murine IL-6 release measured by ELISA in WT, *Caspase 8/Ripk3* DKO, and *Caspase 1/8/11/Ripk3* KO muDCs under the indicated conditions, confirming caspase-independent TLR-driven activation. **(G)** Murine IL-6 release measured by ELISA in WT and *Caspase 1/11* DKO muDCs under the indicated conditions. **(H)** Cell death (% LDH release) in WT and *Caspase 1/11* DKO muDCs under the indicated conditions. **(I)** Immunoblot analysis of NLRP3, caspase-1 (pro-form and p20), caspase-8 (pro-form, p43/41, p18), GSDMD (pro-form and p30), IL-1β (pro-form and p17), and β-actin in WT and *Caspase 8/Ripk3* DKO muDCs stimulated with curli (10, 2.5, 0.0625 µg/mL), L+N, C+N, LPS, or nigericin alone. Data represent mean ± SEM from at least three independent biological replicates. Statistical significance determined by unpaired t-tests with Šídák’s correction or two-way ANOVA. *p < 0.05, **p < 0.01, ***p < 0.001, ****p < 0.0001; ns, not significant.

**Figure S5.**
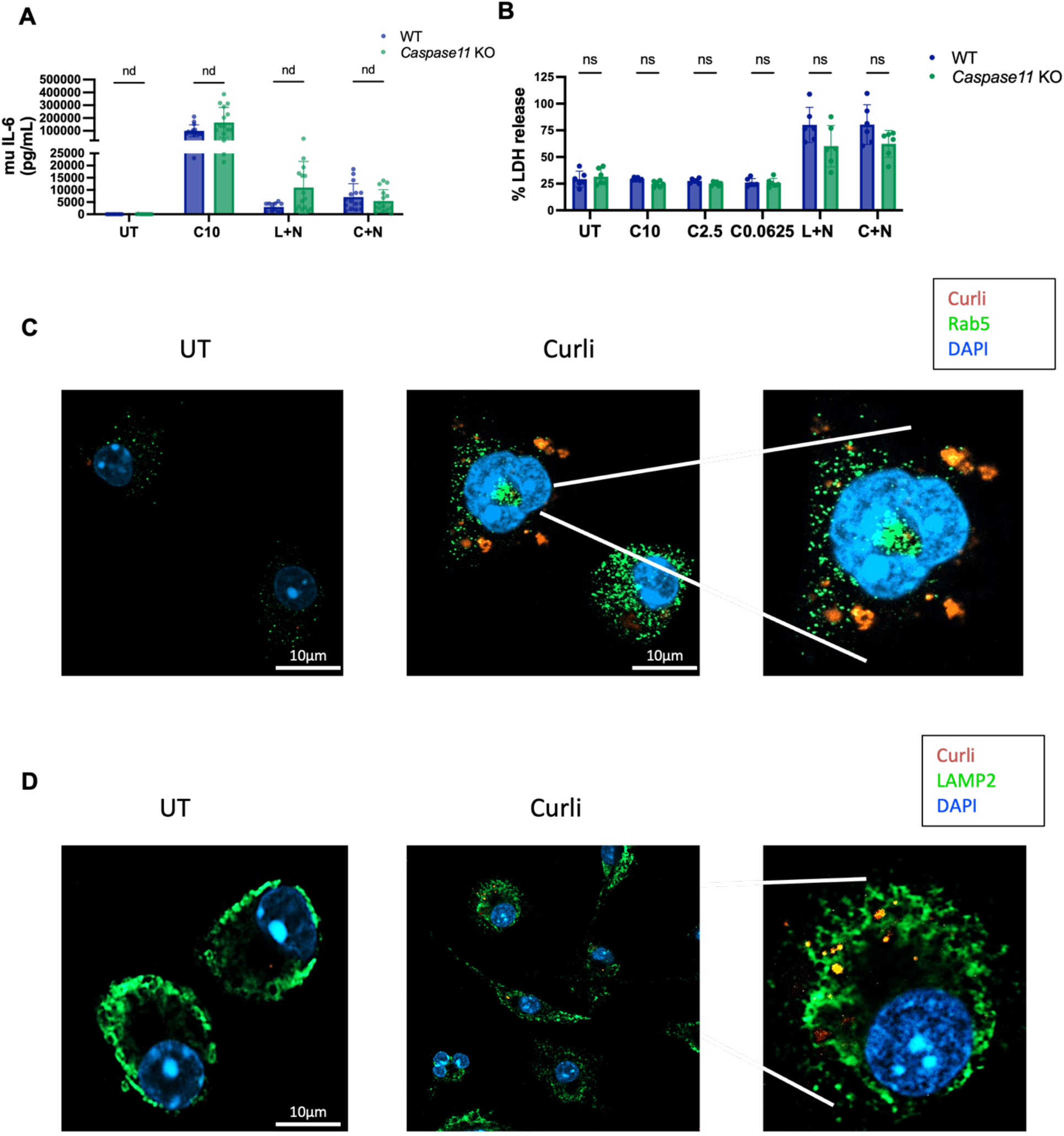
Curli induces IL-6 independent of caspase-11 and traffics through the endolysosomal pathway. **(A)** Murine IL-6 release by ELISA in WT (blue) and *Caspase 11* KO (green) muDCs stimulated with curli (C10), L+N, or C+N, demonstrating caspase-11-independent TLR-driven DC activation. **(B)** Cell death (% LDH release) in WT (blue) and *Caspase 11* KO (green) muDCs stimulated with curli at the indicated concentrations, L+N, or C+N. **(C)** Representative confocal immunofluorescence images of muDCs left untreated or stimulated with Congo Red-labeled curli (orange), co-stained for the early endosomal marker Rab5 (green) and nuclei (blue, DAPI). White lines indicate magnified insets showing co-localization of curli within Rab5-positive early endosomes. Scale bars, 10 µm. **(D)** Representative confocal immunofluorescence images of muDCs left untreated or stimulated with Congo Red-labeled curli (red), co-stained for the lysosomal marker LAMP2 (green) and nuclei (blue, DAPI). White lines indicate magnified insets showing co-localization of curli with LAMP2-positive lysosomes. Yellow puncta in merged images indicate regions of curli-LAMP2 co-localization. Scale bars, 10 µm. Data represent mean ± SEM from at least three independent biological replicates. Statistical significance determined by unpaired t-tests with Šídák’s correction. ns, not significant.

**Figure S6.**
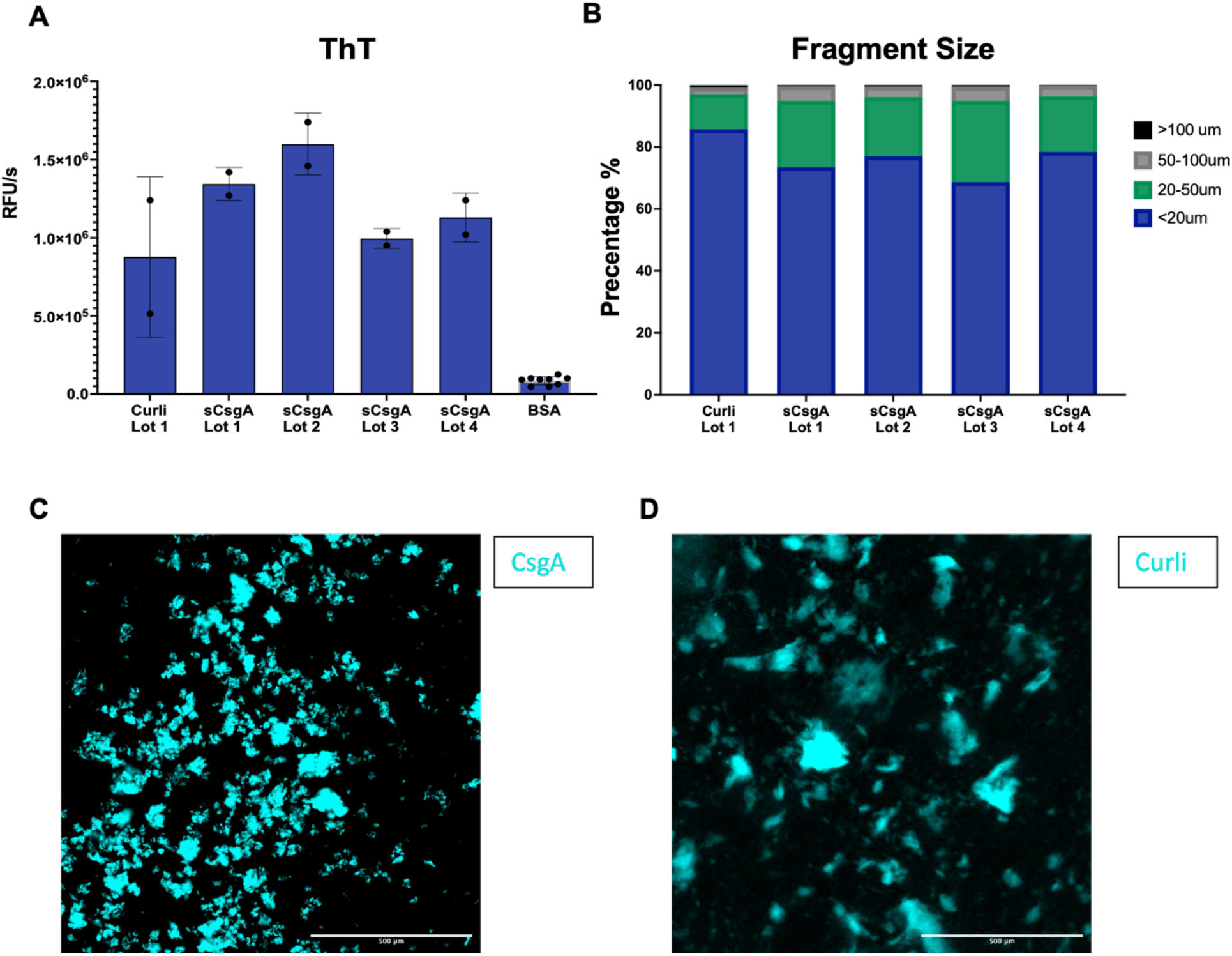
Quality control characterization of naturally purified curli and recombinant fibrillized CsgA preparations. Quality control characterization comparing naturally purified curli (Curli_Lot_1) and four independent lots of recombinant fibrillized CsgA (sCsgA_Lot_1–4), and BSA as negative control. We quantified **(A)** amyloid content assessed by Thioflavin T (ThT) fluorescence (RFU at 10 µg/mL) and **(B)** fibril size distribution (% particles by size range in nm) measured by dynamic light scattering. **(C-D)** Example of fluorescence microscopy of sCsgA_Lot_1 **(C)** and curli **(D)** stained with ThT.

**Figure S7.**
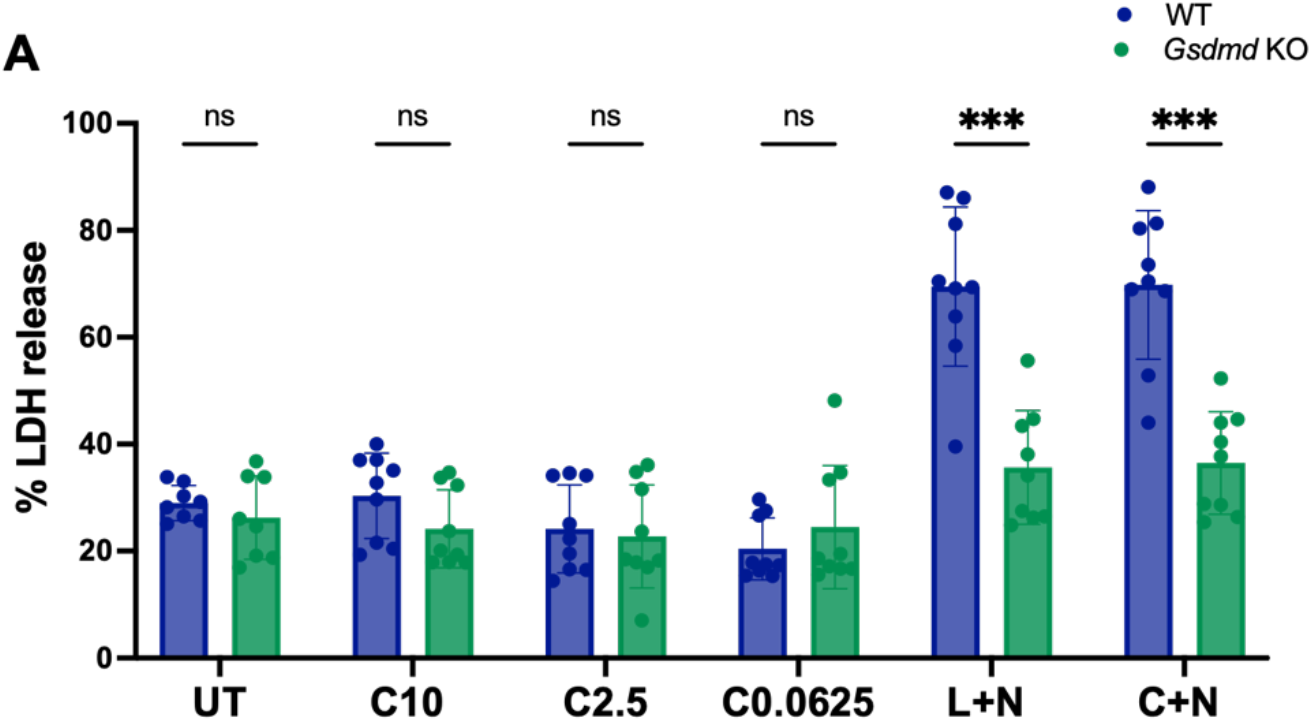
GSDMD deficiency does not affect muDC viability after curli stimulation. **(A)** Cell death (% LDH release) in WT (blue) and *Gsdmd* KO (green) muDCs stimulated with curli (C10, C2.5, C0.0625 µg/mL), L+N, or C+N, demonstrating that GSDMD-dependent lytic cell death is absent in curli-treated conditions but required for pyroptosis induced by L+N and C+N. Data represent mean ± SEM from at least three independent biological replicates. Statistical significance determined by unpaired t-tests with Šídák’s correction between WT and *Gsdmd* KO within each stimulation group. *p < 0.05, **p < 0.01, ***p < 0.001, ****p < 0.0001; ns, not significant.

**Figure S8.**
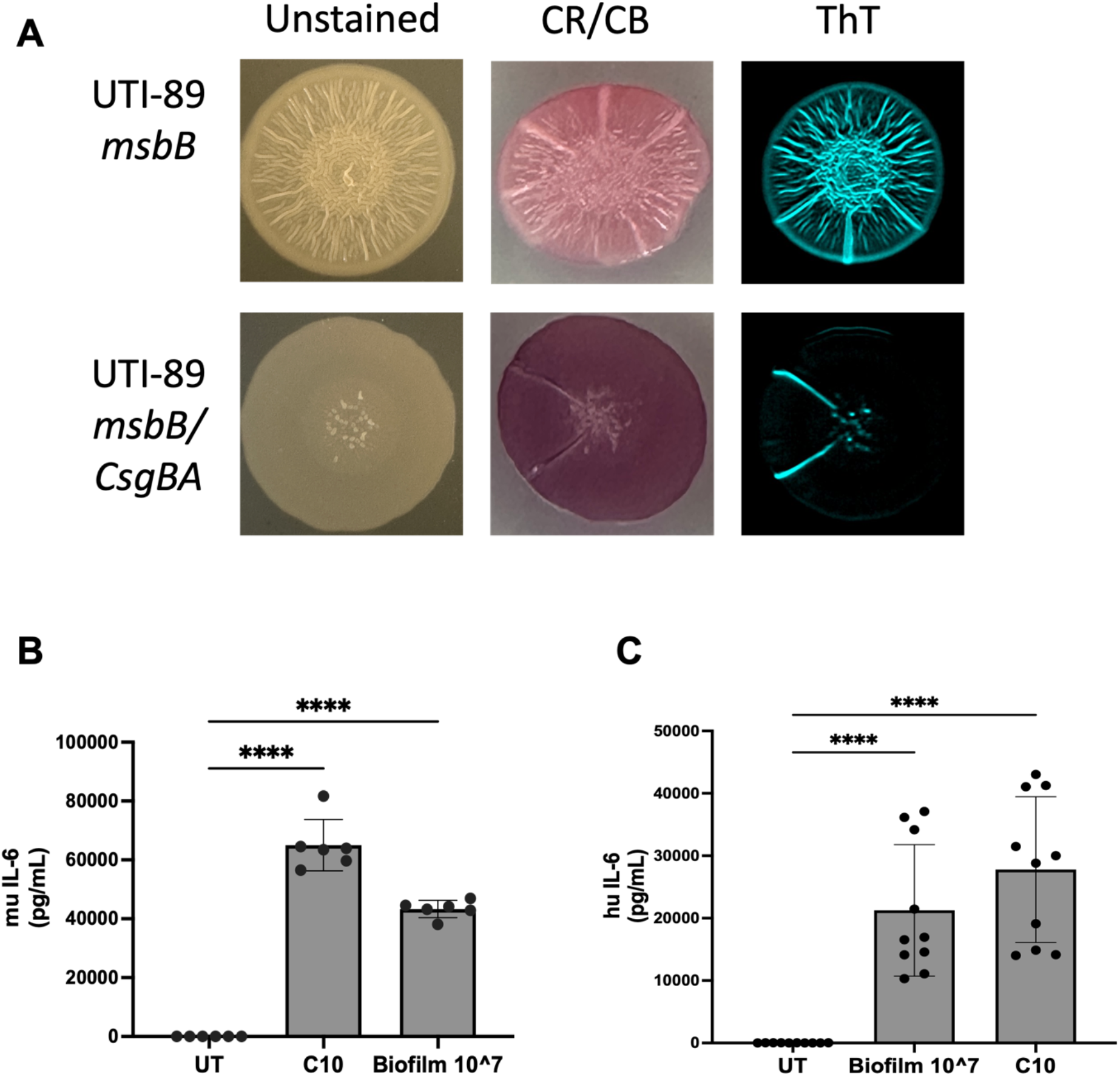
Biofilm induces IL-6 production in murine and human DCs. **(A)** Representative RGB (*left*), Congo Red-Coomassie Blue (CR-CB)(*center*) and Thioflavin (ThT) staining (*right*) images of *E. coli* UTI89-*msbB* (curli-producing, upper panels) and UTI89-*msbB/csgBA* (curli-deficient, lower panels) biofilms grown on YESCA agar for 48 hr at 28°C. Congo Red staining indicates the presence of curli amyloid fibrils in curli-producing strains, which together with Coomassie Blue display as bright pink/red staining and a ridged colony biofilm morphology, whereas curli-deficient colonies show smooth surfaces with dark purple staining from Coomassie Blue. (*right*) ThT fluorescence also indicates the presence of curli amyloid fibrils in the curli-producing *E. coli* UTI89-*msbB* biofilm, and absent in the *msbB/csgBA* one. **(B)** Murine IL-6 and **(C)** human IL-6 were measured by ELISA in WT muDCs stimulated with curli (C10, 10 µg/mL) or UPEC *msbB* biofilm (10^7 eCFU), gentamicin-treated to eliminate viable bacteria prior to DC stimulation. Data represent mean ± SEM from at least three independent biological replicates. Statistical significance determined by one-way ANOVA with Dunnett’s or Holm-Šídák post-hoc test. *p < 0.05, **p < 0.01, ***p < 0.001, ****p < 0.0001; ns, not significant.

